# Characterizing the temporal dynamics and maturation of brain activity during sleep: an EEG microstate study in preterm and full-term infants

**DOI:** 10.1101/2024.03.19.585608

**Authors:** Parvaneh Adibpour, Hala Nasser, Amandine Pedoux, Laurie Devisscher, Nicolas Elbaz, Chloé Ghozland, Elodie Hinnekens, Sara Neumane, Claire Kabdebon, Aline Lefebvre, Anna Kaminska, Lucie Hertz-Pannier, Alice Heneau, Olivier Sibony, Marianne Alison, Catherine Delanoë, Richard Delorme, Marianne Barbu-Roth, Valérie Biran, Jessica Dubois

**Affiliations:** Université Paris Cité, INSERM, NeuroDiderot, F-75019 Paris, France; Université Paris Saclay, CEA, NeuroSpin, UNIACT, F-91191 Gif-sur-Yvette, France; Assistance Publique-Hôpitaux de Paris - APHP, Robert-Debré University Hospital, Department of Physiology – Functional Explorations, F-75019 Paris, France; APHP, Robert-Debré University Hospital, Department of Child and Adolescent Psychiatry, F-75019 Paris, France; APHP, Robert-Debré University Hospital, Department of Pediatric Radiology, F-75019 Paris, France; APHP, Robert-Debré University Hospital, Neonatal Intensive Care Unit, F-75019 Paris, France; Université Paris Cité, CNRS, Integrative Neuroscience and Cognition Center, F-75005 Paris, France; Université Paris Saclay - Université Versailles St Quentin, APHP, Raymond Poincaré University Hospital, Pediatric Physical Medicine and Rehabilitation Department, Garches, France; Université Aix-Marseille, CNRS, Institute of Language, Communication and the Brain, Marseille, France; APHP, Necker-Enfants Malades University Hospital, Department of Clinical Neurophysiology, F-75015 Paris, France; APHP, Robert-Debré University Hospital, Department of Gynecology-Obstetrics, F-75019 Paris, France

**Keywords:** Electroencephalography (EEG), Resting-state, Microstates, Prematurity, Brain development

## Abstract

By interfering with the normal sequence of mechanisms serving the brain maturation, premature birth and related stress can alter perinatal experiences, with potential long-term consequences on a child’s neurodevelopment. The early characterization of brain functioning and maturational changes is thus of critical interest in premature infants who are at high risk of atypical outcomes and could benefit from early diagnosis and dedicated interventions. Using high-density electroencephalography (HD-EEG), we recorded brain activity in extreme and very preterm infants at the equivalent age of pregnancy term (n=43), and longitudinally 2-months later (n=33), compared with full-term born infants (n=14). We characterized the maturation of brain activity by using a dedicated microstate analysis to quantify the spatio-temporal dynamics of the spontaneous transient network activity while controlling for vigilance states. The comparison of premature and full-term infants first showed slower dynamics as well as altered spatio-temporal properties of brain activity in preterm infants. Maturation of functional networks between term-equivalent age and 2 months later in preterms was linked to the emergence of faster dynamics, manifested in part by shorter duration of microstates, as well as an evolution in the spatial organization of the dominant microstates. The inter-individual differences in the temporal dynamics of brain activity at term-equivalent age were further impacted by sex (with slower microstate dynamics in boys) and by gestational age at birth for some microstate dynamics but not by other considered risk factors. This study highlights the potential of the microstate approach to reveal maturational properties of the emerging brain network activity in premature infants.

## 1. Introduction

The last trimester of pregnancy is a critical phase of brain development, with vast growth of long-distance axonal connectivity, onset of myelination in white matter regions, rapid expansion of the cortex, synaptogenesis, and dendritic formation (Kostović & Judaš, 2015). Pre- and peri-natal insults such as preterm birth may impair the progression of these mechanisms, and potentially disrupt the trajectory of neurodevelopment, interfering with post-natal acquisitions and altering long-term cognitive, motor and behavioral outcomes (Arpi & Ferrari., 2013; Bhutta et al., 2002). In particular, infants born before 32 weeks of gestational age (GA) are at a higher risk of adverse neurodevelopmental outcomes (Marlow et al., 2005; Pierrat et al., 2017; 2021).

Magnetic Resonance Imaging (MRI) investigations have associated prematurity with alterations in the brain anatomical maturation by comparing preterm infants at term-equivalent age (TEA) with full-term neonates. These studies have described differences at the macro and microstructural scales, in cortical growth (Ajayi-Obe et al., 2000; Thompson et al., 2019), gyrification (Dubois et al., 2019; Engelhardt et al., 2015), microstructural organization (Bouyssi-Kobar et al., 2018; Dimitrova et al., 2021; Gondova et al., 2023), and white matter connectivity (Ball et al., 2014; Neumane et al., 2022), with differences that could persist into childhood (e.g., Kelly et al., 2023).

However, the functional correlates of such anatomical alterations are still poorly understood despite the common impairments observed at the behavioral level in children born preterm. In the past decade, functional MRI (fMRI) has allowed identifying precursors of spatially organized functional brain networks in the resting-state activity of preterms, and has shown reduced maturation in terms of long-range connectivity compared to full-terms (Doria et al., 2010) and alterations of network-level properties (Fenn-Moltu et al., 2023) that persist until TEA (Gozdas et al., 2018) and childhood (Wehrle et al., 2018). Yet, understanding how these alterations affect the efficiency of brain networks for processing environmental information requires investigating the fast dynamics of the brain activity that go beyond the temporal resolution of the Blood Oxygenation Level Dependent (BOLD) effect captured by fMRI.

While limited in spatial resolution, electroencephalography (EEG) can complement this picture by offering insights into the fast dynamics of the brain activity and by being easily implementable in vulnerable infants. Previous EEG investigations of preterms have indeed identified differences in the occurrence of discontinuous high-amplitude bursts (Whitehead et al., 2018), activity synchronization (Yrjölä et al., 2022), spectral density (Guyer et al., 2019), and in the latency (Kaminska et al., 2017) and scalp distribution of sensory-evoked responses (Maître et al., 2017; Whitehead et al., 2019), with aspects of these alterations persisting into TEA and beyond (Guyer et al., 2019). These investigations indicate that several dimensions of the brain activity such as temporal, scalp distribution, amplitude or signal strength can be impacted by prematurity. Moreover, when investigating the brain spontaneous activity, most methods (i.e. spectral analysis, functional connectivity) study the brain dynamics over several seconds, limiting their interpretation of the temporal properties at a finer scale. Similarly, EEG clinical evaluations involve assessments of sleep structure and maturational figures over relatively long timescales (Dereymaeker et al., 2017), while the brain activity experiences fast changes in its functional organization (Baker et al., 2014), supporting development and learning during sleep (Li et al., 2017). These encourage the use of methods that capture multidimensional information about the brain activity while being sensitive to the fine temporal aspects of the brain dynamics, in order to provide reliable markers of functional alterations in preterm infants.

Microstate analysis aims at identifying short time-segments within the continuous recordings, where the scalp topography (i.e., the distribution of electric potentials originating from the brain activity over the scalp) remains quasi-stable before transitioning into a new scalp topography (Lehmann et al., 1987; Michel & Koenig, 2018). Each of these transient topographies is defined as a microstate (MS), representing the spontaneously occurring transient network activity. Microstates are thus sensitive to both temporal and scalp topographical dynamics of the brain activity.

Recently, the interest in using the microstate approach in infants has emerged, while eliciting various methodological questions (Bagdasarov et al., 2024; Brown et al., 2024; Khazaei et al., 2021). But studies in neonates are rare, focusing on the impact of sleep states on brain dynamics (Khazaei et al., 2021) and on habituation mechanisms to pain (Rupawala et al., 2023), with recent reports indicating microstate dynamics change in the preterm period towards the term-age (Hermans et al., 2023). Yet, the period of the first months after term birth remains unexplored with microstate approach so far, whereas the acceleration of brain functional dynamics and the evolution of network activity topographies are expected to result from the intense progression of maturational processes (e.g. myelination and growth of short cortico-cortical connections) (Kostović & Judaš, 2015). High-density EEG (HD-EEG) recordings might then benefit the characterization of spatio-temporal properties of the brain activity, to better reflect the anatomo-functional changes in the early developmental period. In addition to investigating brain dynamics in the first months after term birth, the potential impact of perinatal risk factors such as prematurity on microstate-related characteristics still needs to be understood. This is of particular importance, given the early onset and heterogeneity of cerebral atypicalities (Dimitrova et al., 2020), requiring the investigation of developmental trajectories through longitudinal assessments.

In this study, we thus aimed to characterize the functional maturation of brain activity dynamics in high-risk preterm infants using HD-EEG recordings. We used the microstate approach to study the maturation of neurophysiological activity between two time-points during development: at term-equivalent age and two months later. First, we compared preterm and full-term born infants in order to identify how the spatio-temporal properties of brain dynamics are impacted by prematurity, while controlling for vigilance states and focusing on sleep states in the main analyses. Next, we implemented a dedicated approach to reliably characterize and compare the organization of spontaneous brain activity in preterms across the two ages. Then, we evaluated how the metrics describing the dynamics of such a functional organization mature over two months after TEA, and we investigated to which extent the inter-individual variability in these dynamics are related to perinatal clinical factors. We hypothesized that spatio-temporal properties of microstates can capture functional aspects of brain activity maturation, which may be sensitive to prematurity and its related risk factors. In particular, we expected an acceleration of the temporal features describing microstates with maturation, which in turn predicted: 1) longer microstate duration (i.e. slower brain dynamics) in preterms vs full-terms, 2) longer microstate duration in infants with higher degrees of perinatal risk factors, 3) shortening of microstate duration from the term-equivalent age to two months later. Since asynchronous maturation of the different brain networks creates distinct time windows of vulnerability across cerebral networks (Neumane et al., 2022; Wu et al., 2017), we did not expect all microstates to be impacted similarly by prematurity.

## 2. Methods

The study design and procedures (protocol DEVine, CEA 100 054) was reviewed and approved by the ethics committee (Comité de Protection des Personnes, CPP Ile de France 3). Parents were informed about the study and gave written consent to participate with their babies. The study involved newborns admitted to the Robert-Debré Children’s Hospital (Paris, France), who were born preterm and followed in the Neonatal Intensive Care Unit, or born full-term in the Maternity.

### 2.1. Participants

This study considered preterm infants born before 32w GA who were due to have a clinical MRI exam at TEA (because of GA lower than 28w or clinical/neurological suspicions) and HD-EEG recordings for research purposes. The MRI exam (obtained in all except one infant) allowed us to measure the Kidokoro score, summarizing the brain development and regional abnormalities (Kidokoro et al., 2013). All infants except one showed a Kidokoro total score (for the whole brain) that was considered as "normal" (score ≤ 3) or related to “mild” abnormalities (4 ≤ score < 8). To favor a relatively homogeneous group, these two infants with no MRI or moderate/severe brain injury were not included in this EEG study.

The preterm cohort then included a group of 43 extreme (born before 28 weeks) and very preterm (born between 28 and 32 weeks) infants (mean GA at birth: 27.1±1.7 weeks, range: [24.1-30.9] weeks), as defined by World Health Organization guidelines (World Health Organization, 2023). Infants were tested longitudinally at two time points: first, at the term-equivalent age (0 months Corrected Age – 0mCA : 41±0.8 weeks of post-menstrual age – w PMA, range: [38.4-42.3] weeks; 20 females, 23 males), and next at 2 months corrected age (2mCA: 50.5±0.7w PMA, range: [49.1-52] weeks; 16 females, 17 males, leading to 33 infants at 2mCA since 10 did not return for the second exam).

Table 1 summarizes the neonatal characteristics of the preterm infants (see Supplementary information, SI Table 1 for the details of individual characteristics). Based on previous studies, we selected and collected data on the clinical factors supposed to play a key role on the neurodevelopment of premature neonates and known as risk factors of adverse outcomes, including GA at birth (considering three GA groups: GA1: 24 weeks+0 day ≤GA≤ 26 weeks+0 day / GA2: 26 weeks+1 day ≤GA≤ 28 weeks+0 day / GA3: 28 weeks+1 day ≤GA≤ 32 weeks+0day), sex, birth weight (correcting for GA and considering values below the 10^th^ percentile as small weight for GA) as well as non-neurological complications during the NICU period: chronic lung disease, need for invasive mechanical ventilation (oxygen therapy) lasting strictly more than 1 day, need for parenteral nutrition longer than 3 weeks, necrotizing enterocolitis, and experiencing sepsis (Brouwer et al., 2017; Neumane et al., 2022). Necrotizing enterocolitis was confirmed by the neonatal clinical team and defined as the presence of clinical evidence meeting the modified criteria for NEC Bell’s stage II, associated with radiologic pneumatosis intestinalis, or stage III determined as definitive intestinal necrosis seen at surgery or autopsy (Walsh & Kliegman 1986). Sepsis was identified in early and late onset cases on the basis of positive blood culture and/or cerebrospinal fluid culture to a pathogen (Sikias et al.,2023; Wynn et al., 2014; US Centers for Disease Control and Prevention, 2024). From these last 5 clinical factors of non-neurological complications, we derived 1) a binary factor of neonatal morbidities associated with prematurity (as in Neumane et al., 2022), to summarize the presence of at least one of them (or the absence of all five), and 2) a continuous ratio score ranging from 0 (no clinical factor) to 1 (all 5 factors) and with 0.2 steps, to summarize how many of the five factors were present in each infant.

**Table 1.**
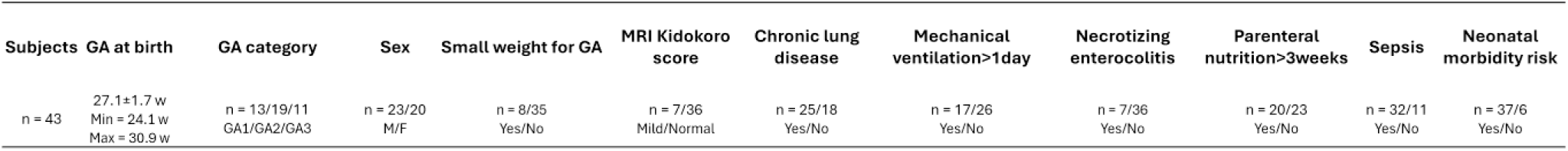
Summary of neonatal characteristics of the infants at birth. Numbers of infants are described over groups of GA at birth (GA1/GA2/GA3), sex (M/F), binarized risk of birth weight indicating small for gestational age (yes/no), binarized MRI Kidokoro score (mild/normal), as well as information regarding five categories of non-neurological complications at NICU. These complications were considered for chronic lung disease (yes/no), use of invasive mechanical ventilation for more than 1 day (yes/no), necrotizing enterocolitis (yes/no), parenteral nutrition for more than 3 weeks (yes/no) and sepsis (yes/no), which were then summarized as one single binarized score of neonatal morbidity factor (yes: at least one of the five identified non-neurological complications/no: none). See Supplementary Table 1 for the report of individual characteristics.

The full-term cohort included a group of 14 infants (mean GA at birth: 40.1±1.1 weeks, range: [37.6-41.6] weeks) who were tested first a few days/weeks after birth (mean age at test : 41.6±1.1w PMA, range: [39.7-43.3] weeks; 7 females, 7 males) and 2 months later (mean age at test : 50.1±0.9w PMA, range: [48.9-51.9] weeks; 7 females, 6 males, leading to 13 infants at 2mCA since 1 infant did not return for the second exam).

### 2.2. EEG recording

High-density 128-channel electroencephalography (HD-EEG, Magstim EGI, Eugene, USA) were recorded on average for approximately 10 minutes during rest, while infants were staying calm in a baby cot, or occasionally in the parent’s arms when it was easier to keep the infant rested. Sampling rate of the recordings was 1000 Hz. A camera recorded the infant’s behavior. The experimenters also inspected infants’ behavior and manually marked the approximate awake/sleep state of infants (based on sustained eye opening or movement) during the recording, in order to allow experts to score sleep stages after the exams (see section 2.4).

### 2.3. Preprocessing

EEG recordings were first bandpass filtered between [0.2-45] Hz, similar to the microstate study in full-term neonates (Khazaei et al.,2021). Note that the high-pass threshold was set slightly higher to better remove low-frequency artifacts. Preprocessing was then performed using an automated pipeline for infants’ continuous EEG (APICE) (Flo et al., 2022). In brief, preprocessing consisted of detecting artifacts on the continuous data where “bad times” and “bad channels” were first defined. Bad times were identified as those with more than 30% of the rejected channels and lasting at least 100ms. Bad channels were those presenting artifacts during more than 30% of the good times. Artifacts were corrected using target principal component analysis on segments shorter than 100ms and spatial spherical spline to interpolate bad channels. This resulted in %0.8±0.6 [%0.1 %1.9] corrected data using target principal component analysis and 10±4.4 [5-19] interpolated channels. Finally, artifacts were detected again, and bad times and channels were re-defined. After applying this automated pipeline, recordings were also visually inspected after this step, and obvious remaining artifacts were marked manually. At the end of this step, segments shorter than 8 seconds were rejected from further analyses. All EEG recordings were down-sampled to 250 Hz and re-referenced to the common-average montage before further processing.

### 2.4. Sleep scoring

For assessing the vigilance state, recordings were first transformed into a low-density bipolar montage as in the standard clinical settings. Different vigilance states were then scored by experts in neonatal electrophysiology. At 0mCA and 2mCA, the recordings were categorized into wakefulness, rapid eye movement (REM) sleep (with low voltage irregular or mixed activity patterns) and non-REM (NREM) sleep (*trace alternant* or high voltage slow activity at 0mCA, and sleep spindles at 2mCA) (Grigg-Damberger., 2016). Wakefulness was distinguished from REM and NREM sleep, based on manual marks from the recording session and camera videos, and considering sustained eye openings and movements. After sleep scoring and preprocessing, we recovered on average 2.5±1.9 [0.7, 7.9] / 6.4±2.5 [1.1, 12.5] / 3.4±2.8 [1.0, 10.9] minutes of recordings for wakefulness / REM sleep / NREM sleep at 0mCA per infant, and similarly 4.1±2.9 [0.8, 11.0] / 5.8±2.7 [1.2, 13.7] / 4.4±2.7 [0.9, 11.5] minutes of recordings at 2mCA (as detailed in section 2.5, not all infants had data in all vigilance states). We only considered comparable vigilance states for age group comparisons. To maintain some homogeneity in terminology, we use REM/NREM labels along the manuscript for both ages, but note that REM/NREM indicate proxies for Active/Quiet sleep at 0mCA (terms that are more conventionally used at this age). The results for REM/NREM states are presented in the main body of the manuscript and those of wakefulness in Supplementary Information, as they involved shorter recording duration and reduced number of infants.

### 2.5. Microstate analysis

Microstate analysis was performed using the Microstate EEGlab toolbox (Delorme & Makeig., 2004; Poulsen et al., 2018), through the following steps: for each subject, 1000 random EEG samples (topographical maps) were taken from the global field power maxima separately for each of the vigilance states. These EEG samples were normalized by the average global field power for each individual to account for the interindividual differences in signal strength (due to non-neuronal differences) and were then concatenated across all subjects and partitioned into a fixed number of microstate classes using modified K-means clustering (Pascual-Marqui et al., 1995) without differentiating opposite polarity topographies (polarity invariance). This allowed identifying the microstate classes (i.e., template microstates) at the group-level per each vigilance state. Clustering was restarted 100 times, and each time the modified k-means was iteratively applied until a convergence threshold of 1e-8 (i.e. relative change in error between iterations) was reached.

For the choice of number of microstate classes, we aimed to maintain consistency with the previous work by Khazaei and colleagues (2021) who reported 7 microstates in full-term neonates (Khazaei et al., 2021). Similarly to their work, we incremented the number of microstate classes between 3 and 15 and estimated how well the template microstates fitted the data by measuring the global explained variance in the data as a function of the number of microstate classes (Figure 1.a). The gain in the global explained variance was less than 1% when more than 7 microstate classes were considered, suggesting that considering more classes would not have a major impact on explaining the variance. To obtain comparable measures to this previous study (Khazaei et al., 2021), we thus selected 7 classes of microstates in further analyses. We also verified that the identified microstates did not demonstrate any features typical of heart or muscle artifacts.

**Figure 1.**
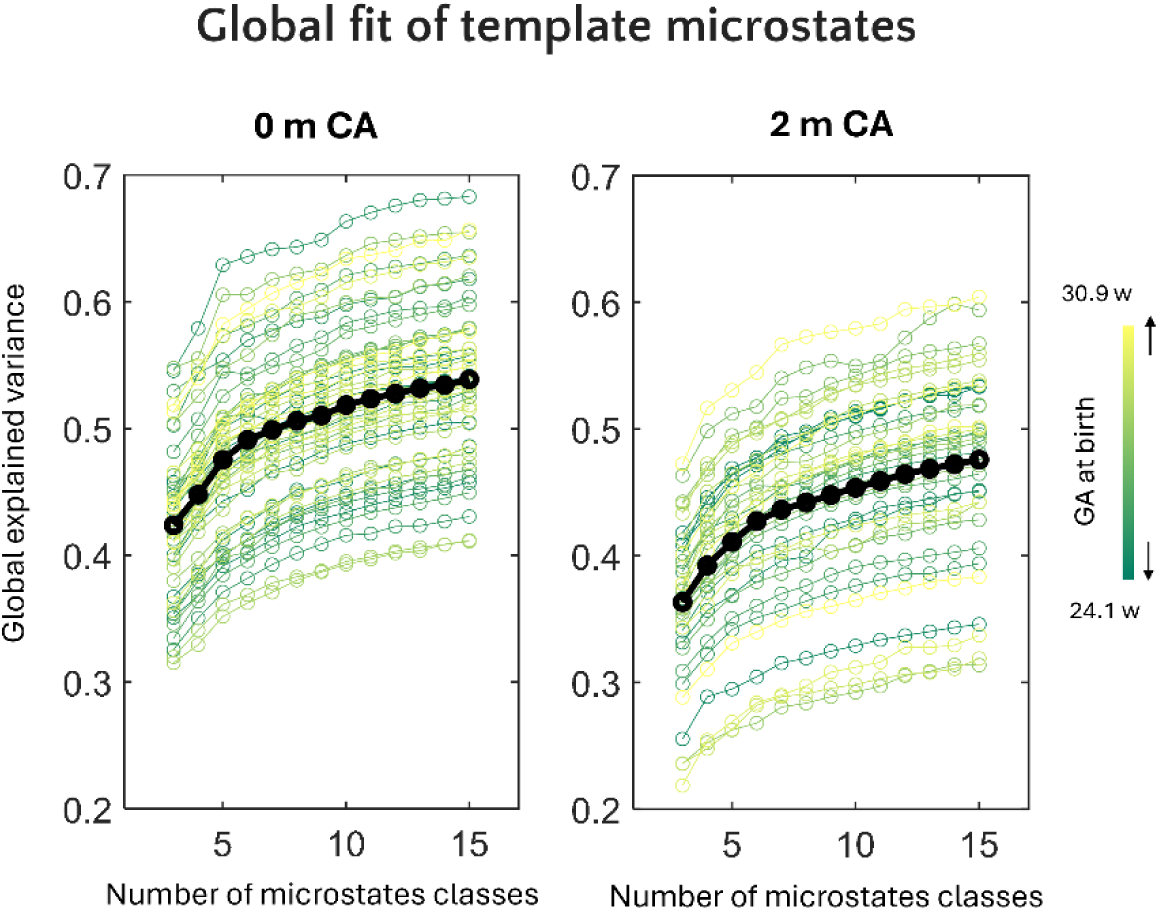
Global fit of the template microstates for different number of microstate classes and different vigilance states. Gain in the global explained variance as a function of the number of microstate classes (3 to 15) for preterm infants at 0 months of corrected age (0mCA) and 2mCA in the overall recordings, without separating vigilance states. Each individual infant is color-coded based on the gestational age (GA) at birth. For a number of microstates higher than 7, the gain in the explained variance dropped below 1%.

The first analysis aimed at comparing microstate characteristics between the preterm and full-term groups in order to identify the potential main effect of prematurity on microstate development. To do so, we focused on identifying the group-level template microstates across both preterm and full-term infants, separately for each vigilance state and each age group (we did not include awake state in this analysis since very few infants in the full-term group had adequate data for this condition). Identifying the template microstates across all infants required accounting for the imbalanced number of subjects in the two groups (more infants in the preterm than the full-term group), in order to avoid biasing the measures to one group. This was achieved by creating five subsets of preterms, where each subset had the same number of infants as the full-term group (0mCA: n_full-terms_=9 for REM sleep / n_full-terms_=10 for NREM sleep; 2mCA: n_full-terms_=8 for REM sleep / n_full-terms_=9 for NREM sleep), including infants from various GA at birth and allowing repetitions across subsets while ensuring that each infant appeared at least in one subset. We identified the 7 template microstates for each of the five preterm subsets completed by the full-term neonates (for each subset: 0mCA: n_full-terms+preterms_ =18 for REM sleep / n_full-terms+preterms_ =20 for NREM sleep; 2mCA: n_full-terms+preterms_ =16 for REM sleep / n_full-terms+preterms_ =18 for NREM sleep). The final set of template microstates, for each of the four conditions (age x sleep stage), was obtained by averaging the template microstates across the five corresponding subsets.

In a second step, analyses focused on the whole preterm group alone in order to evaluate the impact of clinical factors related to prematurity as well as maturational changes between 0 and 2mCA in preterms. The 7 template microstates were identified at the preterm group-level, separately for each vigilance state and each age (0mCA: n=17 for awake / n=41 for REM sleep / n=21 for NREM sleep; 2mCA: n=18 for awake / n =22 for REM sleep / n=23 for NREM sleep).

The procedures above identified group-level template microstates for each of the planned analyses, which were then projected back into each individual infant recording while applying a 30ms temporal smoothing. Back-projecting templates assigned each EEG sample to a microstate class, based on their global topographical dissimilarity invariant to the strength of the signal (Murray et al.,2008). This allowed parsing the continuous EEG recordings of each individual infant into a sequence of microstates, which in turn provided statistical metrics describing the microstate dynamics. These metrics included duration (the average life-time of a microstate when activated), occurrence (the frequency of appearance per second), coverage (the fraction of recording time a given microstate is active), and global explained variance (the similarity of EEG samples to a given microstate class). Only results of the analyses for MS duration are presented in the main manuscript, following our hypothesis that higher maturation is reflected by faster brain dynamics and therefore shorter duration. Results for all other metrics are presented in Supplementary Information, as well as analysis of transition probabilities between different microstates, measuring how frequently one microstate is followed by another microstate.

### 2.6. Statistical analyses

#### 2.6.1. Comparing microstates between the preterm and full-term infants: Impact of prematurity

We first evaluated whether the microstates approach is sensitive enough for capturing the effects of prematurity on the brain activity dynamics. To this end, we compared the metrics of microstate dynamics between preterm and full-term infants separately for each age group and sleep state by conducting Analyses of Variance (ANOVA) with each microstate metric (e.g., duration in the main manuscript) as dependent variable, group (preterm/full-term) as between-subject factor, microstate (MS 1 to 7) as within-subject factor and the two-way interaction group x microstate. This involved conducting 4 ANOVAs per metric (2 age groups, 2 sleep states since wakefulness was not considered).

#### 2.6.2. Comparing microstates between 0mCA and 2mCA: Maturation of brain dynamics in preterms

We assessed how microstates change with maturation between 0 and 2mCA, focusing on the preterm group for three reasons: 1) the results from previous analysis suggested group differences, 2) the main scope of this study focused on prematurity, and 3) the reduced number of full-term infants limited analyses on this group alone. The pipeline implemented to compare microstates characteristics at 0 and 2mCA in preterms is presented in Figure 2 and detailed here. We first compared the two ages at the level of template microstates (0 vs 2mCA) and by measuring their similarity in terms of spatial distribution (Figure 2.a, b). This was achieved by computing the spatial correlation, i.e., absolute Pearson correlation between the vectors of activity values across channels, between the template microstates of the two age groups. This resulted in a correlation matrix, where each value indicated the similarity index between a pair of microstates of the two age groups. To set a threshold for determining similarity of microstates as high or low, we also calculated the spatial correlation between the template microstates within each age group, reasoning that the maximum value of within-group spatial correlations can be considered as the highest spatial similarity between independent states, below which microstates can be considered as independent. Based on this value, we thresholded the two-age-correlation matrix and merged the template microstates with high-similarity by averaging them. This resulted in a set of averaged templates, shared between the two age groups, in addition to non-shared (or non-similar) templates corresponding to the remaining templates with low similarity between the two ages (Figure 2.c). As indicated in the results section, this analysis identified microstates shared between two age groups: 5 for REM sleep, 6 for NREM sleep and 5 for wakefulness. When projecting back the template microstates to each individual infant, we used the shared and non-shared template microstates for each age group (Figure 2.c, d).

**Figure 2.**
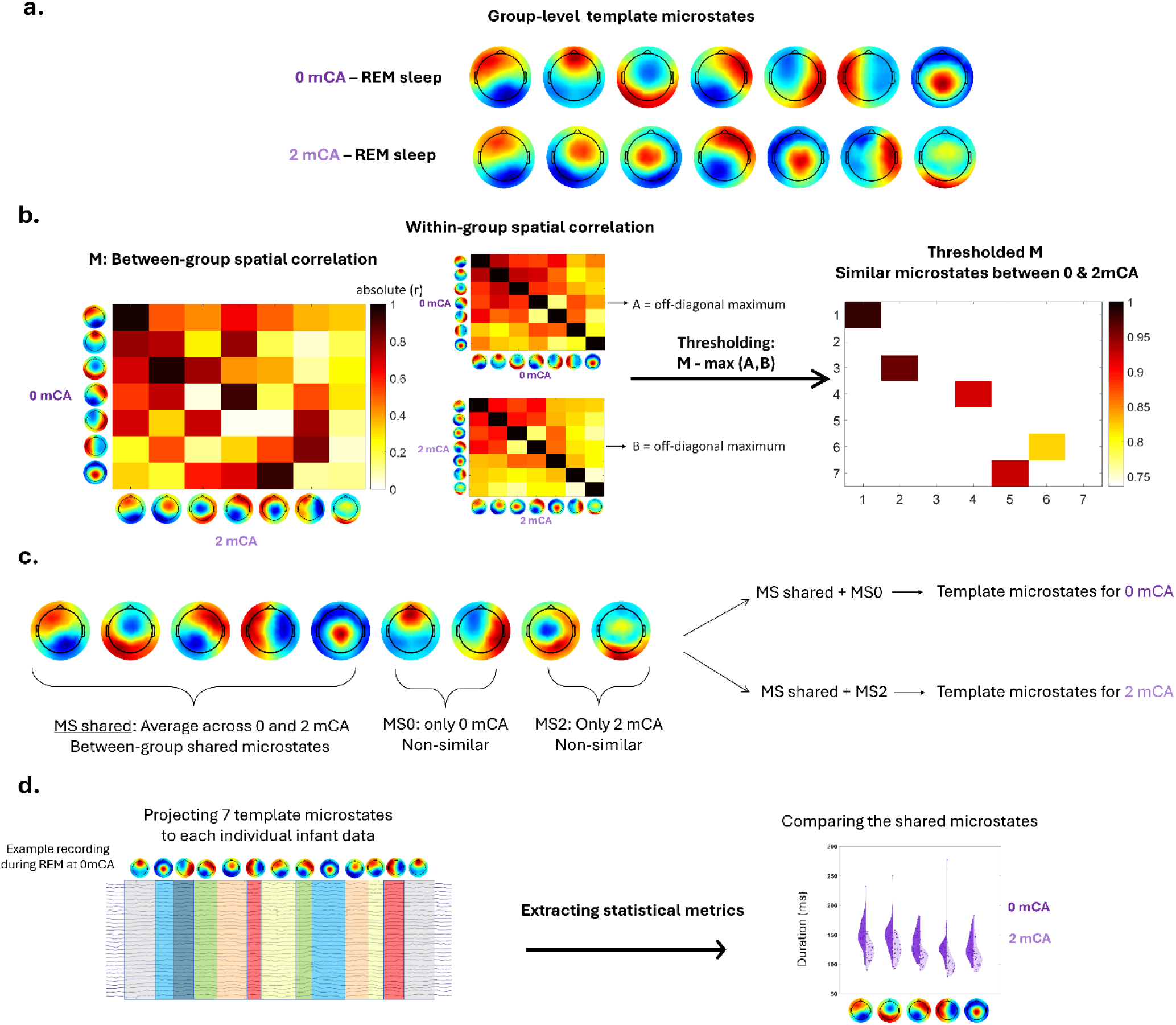
Pipeline comparing microstate characteristics at 0 and 2mCA in preterms in comparable vigilance states. **a.** Group-level template microstates are identified separately for each age group in preterm infants and their head-surface topographical representations are illustrated. **b.** Similarity between group-level microstates of the two age groups is computed using spatial correlation irrespective of polarity of the maps. Similarity between group-level microstates is also computed for each group separately (i.e., within-group) and their maximal values are used to threshold the matrix representing the between-group similarity. **c.** Similar microstates between the two age groups are averaged, constituting the shared microstates (“MS shared”). Non-similar microstates are kept in each age group (0mCA/2mCA → MS0/MS2). The combination of shared and non-similar microstates constituted the final set of 7 template microstates for each age group, among which shared microstates are comparable. **d.** The final template microstates for each age group are back projected to each individual infant recording, illustrating activation of different microstates over time. Microstate metrics are extracted for all the microstates and are compared between age groups only for the shared microstates.

This approach allowed us to compare microstate metrics between the two age groups, focusing on the shared microstates, for each vigilance state. Again, we conducted an ANOVA for each metric (e.g., duration) as dependent variable, age group (0mCA/2mCA) as between-subject factor, microstate as within-subject factor, and the two-way interaction age group x microstate. This involved conducting 3 ANOVAs per metric (3 vigilance states).

Note that few infants had longitudinal data for the same vigilance state: only for REM sleep, an adequate number of infants with longitudinal data (n=21) allowed us to run an additional ANOVA with age group as within-subject factor (i.e., repeated measure) to evaluate whether the results were consistent with the previous analysis with age group as between-subject factor.

As an exploratory analysis, we also investigated whether a structure (i.e. temporal dependency) is observed in the successive activation of microstates, based on transition probabilities (Supplementary Information 3.2).

#### 2.6.3. Relating microstates at 0mCA and their 0 → 2mCA maturation to clinical factors in preterms

To determine how neonatal characteristics at birth and clinical factors during the perinatal period might impact brain maturation, we finally explored the relationships between microstates metrics and the following 5 prematurity risk factors: 1) groups of GA at birth (GA1 / GA2 / GA3); 2) sex (males / females); 3) binarized risk of small weight for gestational age (yes / no); 4) binarized Kidokoro score (mild abnormality / normal); and 5) binarized neonatal morbidity factor (at least one of the 5 identified non-neurological complications / none). To maximize the statistical power and the number of infants, this analysis focused on the conditions of REM sleep at 0mCA (n=41) and at 2mCA (n=21-longitudinally).

A first ANOVA (n=41) assessed the relationship between the MS duration in REM sleep at 0mCA (focusing on microstates for this vigilance state and age) as dependent variable, the 5 risk factors as independent between-subject factors, and microstate (microstate 1 to 7) as independent within-subject factor, as well as specified two-way interactions (microstate x each of the 5 factors, and GA group x sex; all other two-way interactions were not included because they involved few data points). Note that we also performed a similar ANOVA taking into account, not the binary factor of neonatal morbidity, but the 6-level continuous ratio score (no clinical factor to 5 of them), which required considering the averaged MS duration instead of each MS separately, given the limited sample size.

A second ANOVA (n=21) evaluated the relationship between the longitudinal difference in microstate duration in REM sleep between 0mCA and 2mCA (focusing on shared microstates between the two ages) as dependent variable, risk factors (focusing on GA group, sex and binarized neonatal morbidity factor) as independent between-subject factors, microstate (microstate 1 to 5) as independent within-subject factor as well as specific two-way interactions (focusing on microstate x GA group, microstate x sex, and GA group x sex). In this analysis, we did not consider Kidokoro score, the binary factor of small weight for gestational age, as well as other two-ways interactions, because the reduced number of subjects and the reduced inter-individual variability for these factors limited reliable testing.

For all statistical tests, when effects were found to be significant, post hoc tests were carried out using t-tests. And when they involved multiple tests, statistics were corrected for multiple comparisons using the False Discovery Rate (FDR) approach.

## 3. Results

### 3.1. Identifying microstates at different vigilance states

Considering 7 template microstates per age and per sleep states allowed us to describe 0.53/0.59 and 0.45/0.52 of group-level global explained variance for 0mCA and 2mCA groups respectively across REM/NREM sleep. We observed a decrease in global explained variance between 0mCA and 2mCA (p<0.005, Figure 1) and an increase from wakefulness to sleep at both ages (SI Figure 5). The group-level template microstates for different vigilance states and age groups are presented in SI Figure 8.

As mentioned in the methods section, results for MS metrics other than duration as well as all results for awake state, are presented in Supplementary Information (SI.3 and SI.5).

### 3.2. Comparing microstates between preterm and full-term infants: Impact of prematurity

Table 2 (MS duration) and SI Table 2 (other MS metrics) summarize the results of statistical analyses (ANOVA) for the differences between the preterm and full-term groups, at 0 and 2mCA and for each sleep state.

**Table 2.**
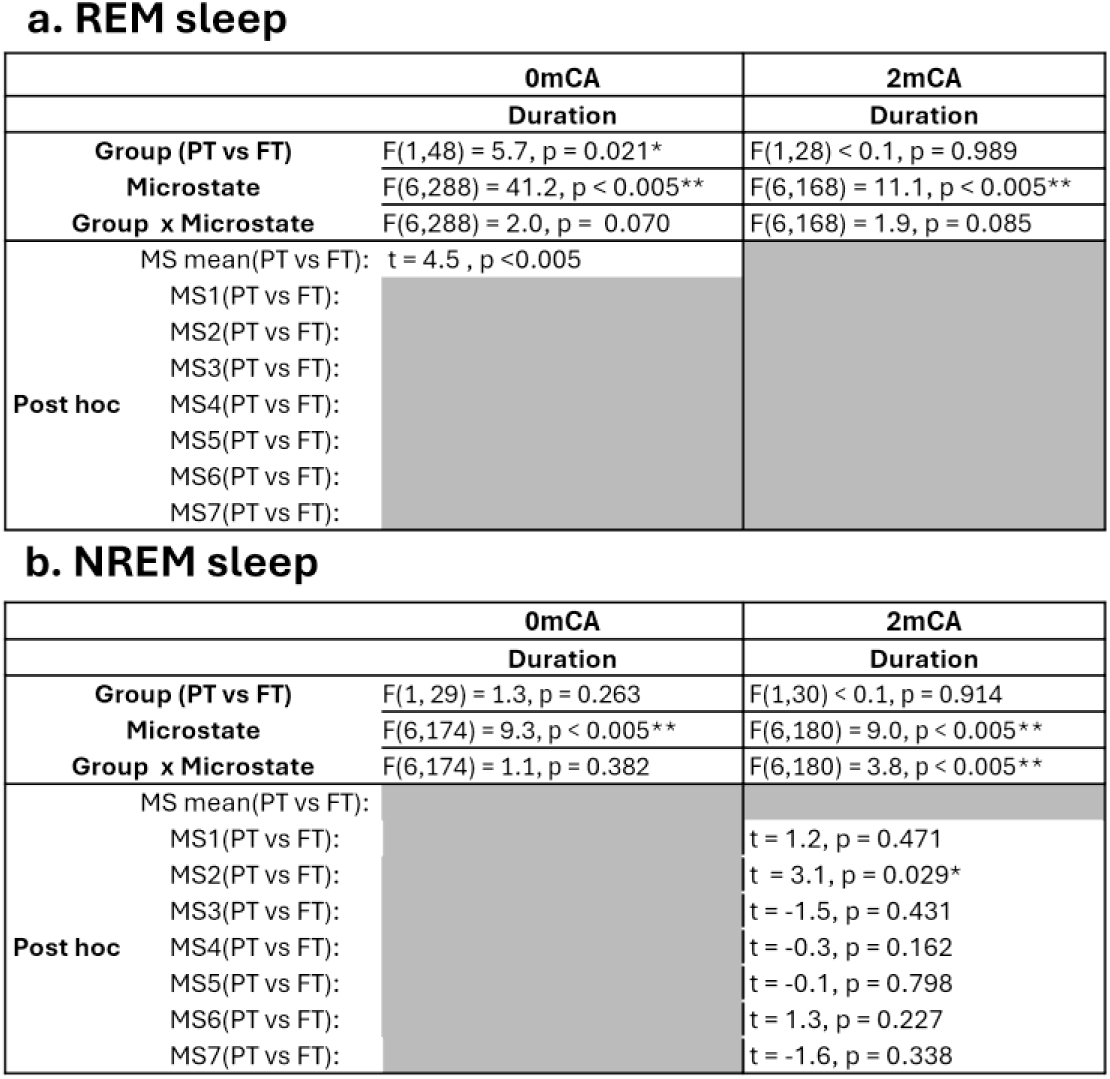
Comparison of microstates (MS) duration between Preterm (PT) and Full-Term (FT) infants for different sleep states at 0 and 2 months of Corrected Age (mCA). Statistical tests were performed for n=41/n=22 preterm vs n=9/n=8 full-term infants at 0mCA/2mCA respectively during REM sleep, as well as for n=21/n=23 preterm vs n=10/n=9 full-term infants at 0mCA/2mCA respectively during NREM sleep. P values were corrected for multiple comparisons with FDR approach for each set of post hoc tests, and significant statistical tests are indicated with asterisks (* p<0.05, ** p<0.005). Comparisons regarding coverage, occurrence and global explained variance are indicated in Supplementary Table 2.

At 0mCA, MS duration in REM sleep showed sensitivity to prematurity: there was a main effect of group (preterm vs full-term, p=0.021) and microstate class (p<0.005) but no interaction between group and microstate class (p>0.05). Post hoc test showed longer duration, averaged across all MS classes, in preterms than full-terms (p<0.005) (Figure 3.a). In NREM sleep, we observed a main effect of microstate class (p<0.005) but no main effect of group or interaction between group and microstate class (all p>0.05) on MS duration (Figure 3.b).

**Figure 3.**
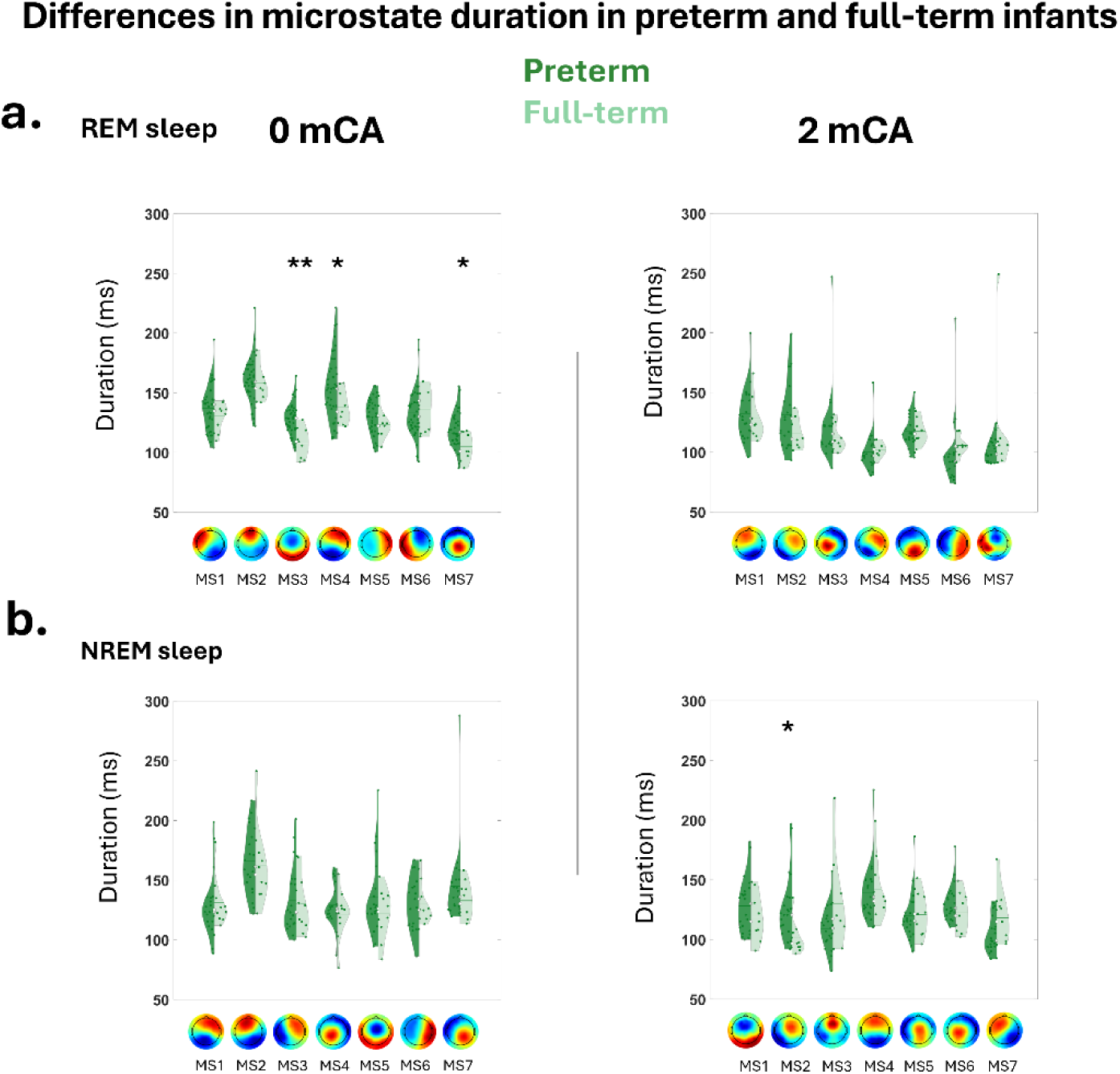
Differences in microstate duration between preterm (dark green) and full-term (light green) infants at 0mCA (left panel) and 2mCA (right panel) during different vigilance states: **a.** REM sleep; **b.** NREM sleep. Asterisks represent significant differences between preterms and full-terms (** p<0.005, *p<0.05). For the other microstate metrics (coverage and occurrence), see SI Figure 1.

At 2mCA, in REM sleep, duration showed a main effect of microstate class (p<0.005) but no significant main effect of the group or interaction between group and microstate class (all p>0.05) (Figure 3.a). In NREM sleep, we observed a main effect of microstate class and an interaction between microstate class x group (all p<0.005), but no main effect of group (p>0.05) (Figure 3.b). Post hoc tests showed longer microstate duration of MS2 in preterms compared to full-terms (p=0.029).

### 3.3. Comparing microstates between 0mCA and 2mCA: Maturation of brain dynamics in preterms

Table 3 (MS duration) and SI Table 3 (other MS metrics) summarize the results of statistical analyses for the changes between 0 and 2mCA, considering different vigilance states and shared MS templates between age groups as detailed in the Methods section and Figure 2.

**Table 3.** Maturation of microstates duration between 0mCA and 2mCA for different sleep states. Statistical comparisons were performed for n=41 at 0mCA vs n=22 at 2mCA preterm infants, during REM and for n=21 0mCA vs n=23 at 2mCA preterm infants, during NREM sleep. The statistical tests in a. and c. were made cross-sectionally and in b. for n=21 infants longitudinally. P values were corrected for multiple comparisons with FDR approach for each set of post hoc tests, and significant statistical tests are indicated with asterisks (* p<0.05, ** p<0.005). Comparisons regarding coverage, occurrence and global explained variance are indicated in Supplementary Table 3.

**a. REM sleep**
| Duration |  |
| --- | --- |
| **Age (0 vs 2mCA)** | F(1,61) = 33.8, p < 0.005 ** |
| **Microstate** | F(4,244) = 14.1, p < 0.005** |
| **Age x Microstate** | F(4,244) = 0.9, p = 0.453 |
| **Post hoc** | MS mean (0 vs 2mCA): t = 5.6, p<0.005 |
|  | MS1(0 vs 2mCA): |
|  | MS2(0 vs 2mCA): |
|  | MS3(0 vs 2mCA): |
|  | MS4(0 vs 2mCA): |
|  | MS5(0 vs 2mCA): |

**b. REM sleep (longitudinal)**
| Duration |  |
| --- | --- |
| **Age (0 vs 2mCA)** | F(1,20) = 46.1, p < 0.005** |
| **Microstate** | F(4,80) = 8.3, p < 0.005** |
| **Age x Microstate** | F(4,100) = 0.6, p = 0.630 |
| **Post hoc** | MS mean (0 vs 2mCA): t = 4.9, p<0.005 |
|  | MS1(0 vs 2mCA): |
|  | MS2(0 vs 2mCA): |
|  | MS3(0 vs 2mCA): |
|  | MS4(0 vs 2mCA): |
|  | MS5(0 vs 2mCA): |

**c. NREM sleep**
| Duration |  |
| --- | --- |
| **Age (0 vs 2mCA)** | F(1,42) = 6.1, p = 0.018 * |
| **Microstate** | F(5,210) = 13.1, p < 0.005** |
| **Age x Microstate** | F(5,210) = 5.9, p < 0.005** |
| **Post hoc** | MS mean (0 vs 2mCA): |
|  | MS1(0 vs 2mCA): t = 1.6, p = 0.186 |
|  | MS2(0 vs 2mCA): t = 3.0, p = 0.015* |
|  | MS3(0 vs 2mCA): t = 0.9, p = 0.389 |
|  | MS4(0 vs 2mCA): t = -2.2, p = 0.066 |
|  | MS5(0 vs 2mCA): t = 4.0, p < 0.005** |
|  | MS6(0 vs 2mCA): t = 1.4, p = 0.202 |

For REM sleep, 5 microstates out of 7 were shared between 0 and 2mCA and therefore included in the ANOVA (Table 3.a). We observed a main effect of age group and microstate class on the MS duration (all p<0.005) but no interaction between age group and microstate class (p>0.05). Post hoc tests indicated shorter microstate durations, averaged across all MSc classes, at 2mCA compared to 0mCA (p<0.005, Figure 4.a). When restricting infants to the subgroup with longitudinal data both at 0 and 2mCA for REM sleep, results were replicated (Table 3.b).

**Figure 4.**
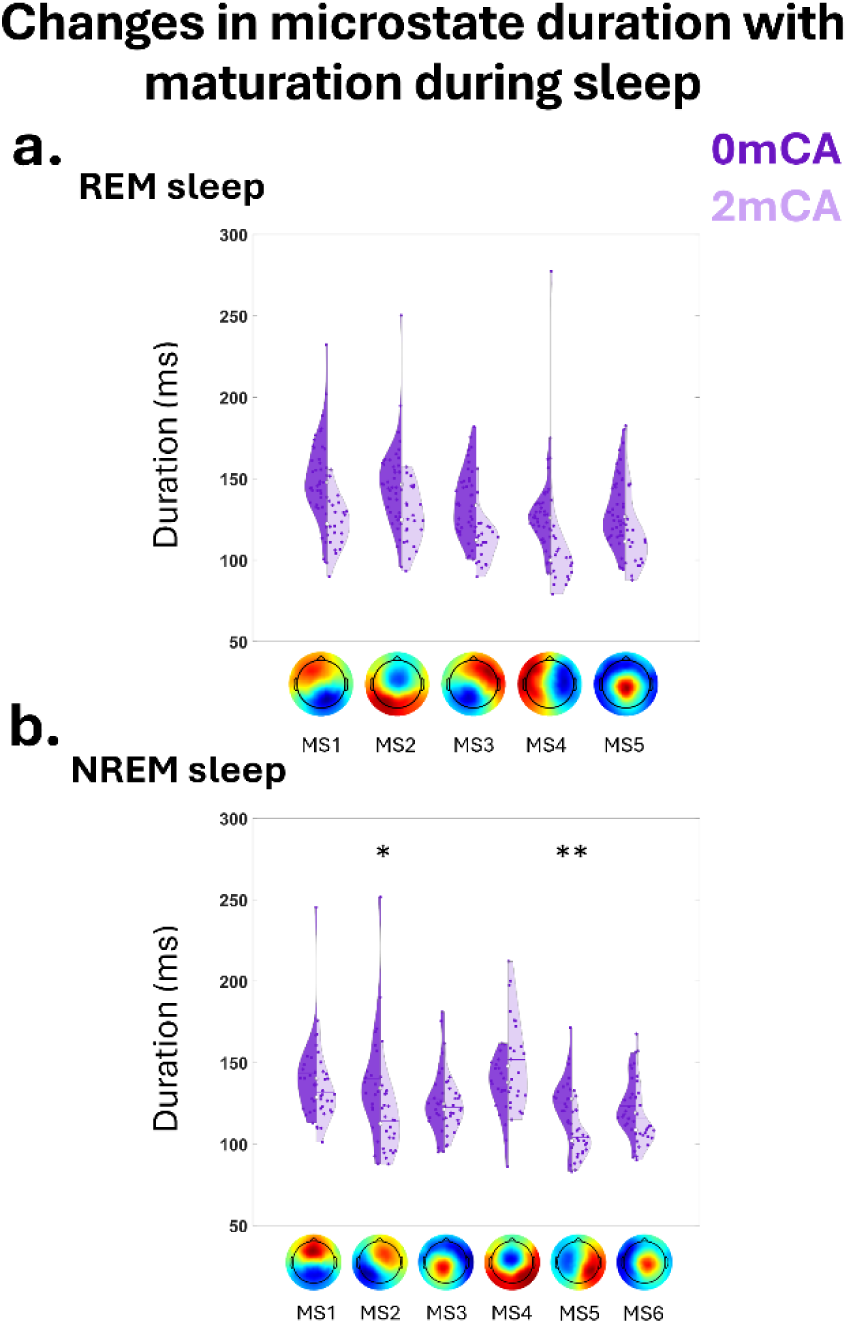
Evolution in microstate duration between 0mCA (dark purple) and 2mCA (light purple) in preterms, during different sleep states: **a**. for the 5 shared microstates identified in REM sleep; **b**. for the 6 shared microstates in NREM sleep. Asterisks represent significant differences between 0mCA and 2mCA infants (** p<0.005, *p<0.05). For the other microstate metrics (coverage, occurrence and global explained variance), see SI Figure 1.

For NREM sleep, 6 microstates out of 7 were shared between 0 and 2mCA and therefore included in the ANOVA (Table 3.c). We observed a main effect of age group (p=0.018) and microstate class (p<0.005) and an interaction between age group and microstate class (p<0.005) on MS duration. Post hoc analyses indicated that MS2 and MS5 showed shorter duration at 2mCA compared to 0mCA (MS2: p=0.015, MS5: p<0.005) (Figure 4.b).

The microstates not shared between the 0 and 2mCA groups suggested differences in the spatial distribution of network activity. In REM sleep, non-shared microstates showed more abundant posterior and central/posterior topographical distributions at 2mCA compared to 0mCA (Figure 2 and SI Figure 8.c.) while in NREM sleep, they showed more abundant presence of anterior/posterior topographical distributions at 2 mCA (SI Figure 8.c.).

For the sake of simplicity, the transition probabilities between microstates are described in Supplementary Information (SI Figure 3).

### 3.4. Relating perinatal factors to microstates at 0mCA and their maturation between 0mCA and 2mCA

Finally, we performed statistical analyses to evaluate the impact of perinatal factors on the microstate duration, first at 0mCA and focusing on REM sleep to allow further longitudinal comparisons at 2mCA. We observed a main effect of sex (p<0.005), microstate class (p<0.005), as well as an interaction between group of GA at birth and microstate class (p=0.016), but no main effect of group of GA at birth, Kidokoro score, birth weight, neonatal morbidity or other considered interactions (all p>0.05) (Figure 5, Table 4). Post hoc tests revealed longer duration, averaged across all MS classes, in males than females (p<0.005), longer duration for GA1 compared to GA2 for two MS, but also for GA3 compared to GA2 for one MS (p<0.05). Complementary analysis considering the averaged MS duration and 4 groups of GA at birth (GA1, GA2, GA3, full-term) confirmed that preterms had longer duration than full-terms (SI Figure 4, SI Table 4).

**Figure 5.**
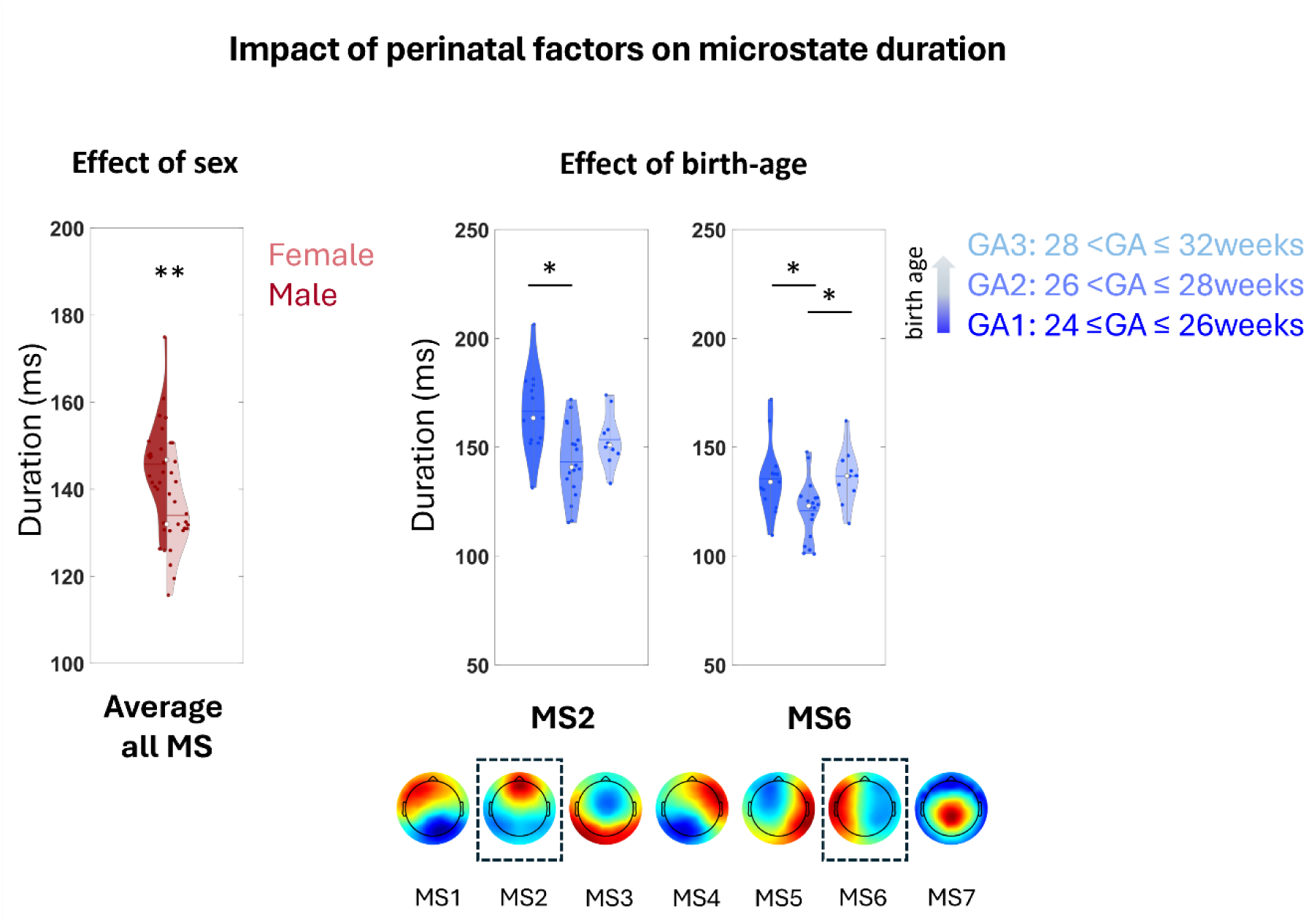
Impact of perinatal factors on microstate duration at 0mCA for infants born preterm. Impact of **a.** Sex, considering the average duration over all MS; **b.** Groups of GA at birth for specific MS of REM sleep. Asterisks represent significant differences between sex and GA groups (** p<0.005, *p<0.05): longer duration in males and in infants with the lowest GA at birth.

**Table 4.** Impact of perinatal factors on microstates at 0mCA and their maturation between 0mCA and 2mCA. The relationships between microstate duration at 0mCA (n = 41, left column) or the longitudinal changes in microstate duration between 0mCA and 2mCA (n=21, right column) and different prematurity risk factors were evaluated for REM sleep activity. These factors included group of GA at birth (GA1/GA2/GA3); sex; binarized risk of birth weight indicating small for GA; MRI Kidokoro score and binarized neonatal morbidity factor summarizing non-neurological complications. At 2mCA, only main effects and interactions with adequate data points were considered. P values were corrected for multiple comparisons with FDR approach for each set of post hoc tests, and significant statistical tests are indicated with asterisks (* p<0.05, ** p<0.005).

|  |  | 0mCA | 0mCA→2mCA |
| --- | --- | --- | --- |
| **GA group** |  | F(2,31) = 2.6, p = 0.090 | F(2,14) = 1.9, p = 0.185 |
| **Microstate** |  | F(6,203) = 12.6, p < 0.005** | F(4,68) = 0.7, p = 0.545 |
| **MRI Score** |  | F(1,31) = 0.5, p = 0.474 |  |
| **Sex** |  | F(1,31) = 15.9, p < 0.005** | F(1,14) = 2.3, p = 0.151 |
| **Small Weight for GA** |  | F(1,31) = 4.0, p = 0.054 |  |
| **Morbidity Risk** |  | F(1,31) < 0.1, p = 0.803 | F(1,14) < 0.1, p = 0.871 |
| **GA group x Sex** |  | F(2,31) < 0.1, p = 0.928 | F(2,14) < 0.1, p = 0.956 |
| **Microstate x GA group** |  | F(12,23) = 2.1, p = 0.016* | F(8,68) = 0.5, p = 0.866 |
| **Microstate x MRI score** |  | F(6,203) = 1.2, p = 0.290 |  |
| **Microstate x Sex** |  | F(6,203) = 1.2, p = 0.321 | F(4,68) = 1.0, p = 0.375 |
| **Microstate x Small Weight for GA** |  | F(6,203) = 0.7, p = 0.619 |  |
| **Microstate x Morbidity Risk** |  | F(6,203) = 0.8, p = 0.536 |  |
| **Post hoc** | MS mean : Female vs Male | t = -3.5, p < 0.005 |  |
|  | MS1: GA1 vs GA2 | t = 0.7, p = 0.734 |  |
|  | MS1: GA1 vs GA3 | t = -0.1, p = 0.900 |  |
|  | MS1: GA2 vs GA3 | t = -0.9, p = 0.734 |  |
|  | MS2: GA1 vs GA2 | t = 3.5, p = 0.005** |  |
|  | MS2: GA1 vs GA3 | t = 2.0, p = 0.074 |  |
|  | MS2: GA2 vs GA3 | t = -1.9, p = 0.074 |  |
|  | MS3: GA1 vs GA2 | t = -0.9, p = 0.816 |  |
|  | MS3: GA1 vs GA3 | t = -0.6, p = 0.816 |  |
|  | MS3: GA2 vs GA3 | t = -0.1, p = 0.937 |  |
|  | MS4: GA1 vs GA2 | t = 1.6, p = 0.249 |  |
|  | MS4: GA1 vs GA3 | t = 1.4, p = 0.249 |  |
|  | MS4: GA2 vs GA3 | t < 0.1, p = 0.982 |  |
|  | MS5: GA1 vs GA2 | t = 1.5, p = 0.223 |  |
|  | MS5: GA1 vs GA3 | t = -0.7, p = 0.488 |  |
|  | MS5: GA2 vs GA3 | t = -2.3, p = 0.095 |  |
|  | MS6: GA1 vs GA2 | t = 2.6, p = 0.024* |  |
|  | MS6: GA1 vs GA3 | t = -0.2, p = 0.867 |  |
|  | MS6: GA2 vs GA3 | t = -3.0, p = 0.021* |  |
|  | MS7: GA1 vs GA2 | t = -0.7, p = 0.751 |  |
|  | MS7: GA1 vs GA3 | t = 0.3, p = 0.751 |  |
|  | MS7: GA2 vs GA3 | t = 0.9, p = 0.751 |  |

For longitudinal changes in MS duration from 0mCA to 2mCA, during REM sleep, we did not observe any significant effect (Table 4; SI Table 4).

## 4. Discussion

In this study, we used an original microstate approach to characterize the fast transient dynamics of EEG activity in preterm and full-term infants while considering different vigilance states. We reported three sets of results showing that this approach is sensitive for capturing functional aspects of brain maturation and dysmaturation. First, by comparing preterms and full-terms, we identified alterations at term-equivalent age that also persisted at 2mCA, indicating the impact of prematurity on early brain activity patterns. Second, comparing EEG dynamics between 0 and 2mCA in preterms with a robust approach allowed us to identify the presence of both similar (i.e., shared between age groups) and unsimilar microstates in all vigilance states. For microstates similar between age groups, we observed a shortening of microstate duration, as well as changes in the coverage, occurrence, global explained variance, and the transition patterns of microstates (Supplementary Information). These highlighted an evolution in the spatio-temporal properties of brain activity with maturation. Third, we showed that the inter-individual differences in the temporal dynamics of microstates in preterms is impacted by gestational age at birth and sex, with longer microstate duration in infants with lower birth age and in males. This confirmed the sensitivity of the implemented approach in capturing the effects of risk factors on early brain activity.

### Brain dynamics are impacted by prematurity and dysmaturation

Comparing preterm and full-term infants indicated that the microstate dynamics, as captured with MS duration and other metrics, are impacted by prematurity, particularly at TEA. These alterations were primarily observed in terms of slower dynamics in preterms (i.e., longer duration and lower occurrence rate of microstates), which may find their roots in less mature brain structures. Such slower dynamics might reflect less advanced or altered myelination of white matter pathways (Neumane et al, 2022), as this mechanism accelerates the conduction of neural impulses and impact the speed of brain responses during development (Dubois et al., 2008), and/or to less complex cortical microstructure with reduced dendritic arborization and synaptogenesis (Dimitrova et al., 2021; Gondova et al., 2023), as this might negatively impact neural activity synchronization (Dupont et al., 2006). These cerebral alterations in preterms at TEA may therefore affect the network-wide activity patterns we measured with the MS approach. Similar slower brain dynamics has also been recently reported in infants with hypoxic-ischemic encephalopathy (Khazaei et al., 2023), confirming that perinatal insults delay the development of network-wide brain activity.

However, direct links between the structural and functional alterations remain to be established, potentially by delineating different brain networks that mature progressively during infancy (Dubois et al., 2014, 2016). In line with this proposition, we also observed that the impact of prematurity differed across microstate classes (interaction group x microstate class) in some conditions (i.e. NREM at 2mCA), which suggests that not all functional networks are impacted in a similar way. This is further in agreement with the well-known asynchronous maturation pattern of the different brain networks (Yakovlev & Lecours, 1967 & Dubois et al., 2014), suggesting that distinct time windows of vulnerability (Wu et al., 2017) lead to differential adverse impacts across cerebral networks (Neumane et al., 2022). All together these findings encourage studying structure-function relations and alterations by considering different networks separately.

Beyond prematurity, we also aimed to evaluate to which extent specific perinatal and clinical factors might impact the development of brain activity dynamics in infants. Previous studies have identified several risk factors for poorer outcomes related to prematurity (Allen et al., 2008; O’Driscoll et al., 2018; Pierrat et al., 2021). Nevertheless, our sample size limited the possibility to test multiple associations, so we focused on the factors that supposedly have a major impact. First, GA at birth appeared to modulate the microstate dynamics and maturation: preterm infants within GA1 group (24w ≤GA≤ 26 w) showed slower dynamics than GA2 group (26w <GA≤ 28w) and full-term infants (Supplementary Information 4.1), in a comparable way as the effect of preterm vs full-term birth. Yet, while our preterm cohort had quite uniform sampling of gestational ages at birth between 24 and 32 weeks, we did not observe a dose-dependent effect of gestational age on microstate duration, as GA3 group (28w <GA≤ 32w) did not show shorter MS duration compared to GA1 and GA2 groups. This might be explained by the heterogeneity of the cohort clinical characteristics across GA groups. In fact, only infants due to have a clinical MRI at TEA were included in our study, either because they were born before 28w GA (GA1 and GA2 groups) and/or because of clinical/neurological suspicions (GA3 group). Thus, contrarily to GA1 and GA2 groups, GA3 group was not an exhaustive representation of preterm infants born between 28 and 32w GA, even though no infant showed moderate or severe brain abnormalities according to MRI Kidokoro score. Importantly, GA at birth had a disparate impact across the different microstates, suggesting that different networks have different time windows of vulnerability over the gestational period (Wu et al., 2017). In future studies, it would be interesting to characterize such an effect by considering GA as a continuous variable in a larger group of infants including some with moderate prematurity (GA at birth between 32 and 37w).

We also evaluated the impact of sex on microstates dynamics and our results of slower dynamics in males seemed consistent with the large clinical and epidemiological evidence of higher vulnerability of males compared to females over this specific period of development (Kelly et al., 2023; Hintz et al., 2006; Perivier et al., 2016). These results further suggested the sensitivity of the microstate approach to highlight some inter-individual variability across infants, that might be interesting to target for individualized interventions. Nevertheless, controlling for the known sex differences in brain size might be an important aspect in future studies to disentangle the effects of sex and brain anatomical characteristics on brain dynamics.

Other clinical risk factors of adverse outcomes related to prematurity were also considered (small birth weight, respiratory, intestinal or infectious complications and brain abnormalities) but none were shown to impact the brain dynamics in our study. Nevertheless, this may be related to the lack of large variability in our cohort of infants and to the limited sample size which constrained us to binarize factors and combine some into a composite morbidity score. It is also possible that some of the assessed clinical factors were not reflective of maturational changes captured with EEG dynamics. In particular, the MRI Kidokoro score is supposed to capture macrostructural abnormalities related to global and regional volumetric growth as well as brain lesions but may not be sensitive to diffuse microstructural alterations impacting the network functional activity.

### Maturation changes the spatio-temporal brain dynamics in preterm infants

Comparing the dynamics of the brain activity between 0m and 2mCA indicated an evolution in terms of distinctive spatial topographies of activity among the dominant microstates, as well as major impact on temporal properties of microstates already developed early on. The presence of a few non-similar MS across groups might further suggest a reorganization of functional network activities during this period of intense maturation in cortical regions and white matter pathways promoting functional specialization of networks (Dubois et al., 2014; Gao et al., 2014; Keunen, Counsell, Benders, 2017). For example, the appearance of a majorly posteriorly-distributed MS at 2mCA in REM sleep might relate to the intense post-natal development of the visual system at these ages (Dubois et al.,2008; 2014).

The observed major shortening of microstate duration between 0mCA and 2mCA is also in agreement with a recent study of preterm infants indicating acceleration of MS dynamics between the preterm and term period [250–300ms at 30w PMA to 150–200ms after 37w PMA] (Hermans et al., 2023). Interestingly, the average duration of the microstates in our study, 140ms at 0mCA and 112ms at 2mCA, is in the intermediate range between the preterm period (Hermans et al., 2023) and childhood with microstate durations reported between 60 to 90ms (Bagdasarov et al., 2022; Koenig et al., 2002; Takarae et al., 2021). This suggests a continuous change in the duration of MS from birth through childhood. The shortening of MS duration, together with the increase in the occurrence rate of MS, indicate an acceleration of the brain dynamics which could reflect the intensive myelination of white matter pathways (Dubois et al., 2008), and/or maturation of cortical microstructure (Dimitrova et al., 2021; Gondova et al., 2023) around TEA and first post-term weeks resulting in an acceleration and better synchronization of neural dynamics. Several developmental changes concerned microstates with an anterior/posterior distribution of the brain activity, a spatial organization also reported to show alterations of functional connectivity patterns in preterms (Omidvarnia et al., 2014; Tokariev et al., 2019). An interesting direction for future investigations would be to address how the functional connectivity patterns identified with EEG relate to the dynamics of microstates and their maturation in early development (see Khazaei et al., 2023).

While maturational changes between 0mcA and 2mCA at REM affected the duration of all MS classes, at NREM sleep these changes concerned some MS classes which might in part reflect the maturation changes in sleep structure and in particular the appearance of sleep spindles in NREM sleep after 43 weeks of post-menstrual age (Grigg-Damberger.,2016).

Besides metrics describing individual microstates, we also explored the transitions between microstates (Supplementary Information 3.2.) which offer insights into the structure of the temporal dependencies in brain dynamics. At both ages, the transitions between the microstates had a non-random structure, with most microstates tending to transition into their ‘favorite’ microstates (one or more than one). The pattern of such transitions evolved with development between 0 and 2mCA, which could suggest a reorganization of the brain functional dynamics, showing an evolution in the continuous shifting between transient network activities. Beyond microstates, dynamic aspects of brain activity have also been described for functional brain networks identified with fMRI at the temporal resolution of the BOLD effect (França et al., 2024), and with functional ultrasound imaging (Baranger et al., 2021), indicating their alterations in preterms before and at TEA compared to full-terms. Here, we could not compare the transition patterns of microstates between the preterm and full-term groups, because of the limited sample size of the latter group.

### Methodological considerations for the characterization of microstate dynamics in infants

First, across all analyses, we ensured that the group-level microstate templates were not biased towards a subgroup of infants. For the preterms vs full-terms comparison, we presumed the microstate templates should be similar between the preterm infants at TEA and full-term neonates, given that the group-level microstates identified in the whole preterm group during sleep were qualitatively similar to those of a previous study in full-term asleep neonates (Khazaei et al., 2021). Yet, to ensure that the initially-defined templates were equally representative of both groups and unbiased towards certain GA at birth, we implemented a strategy iterating over subgroups of preterms with the same number as full-terms and with various GA. Consistently, no correlation between GA and the metric of global explain variance was then found. Similarly, we ensured that comparing microstate metrics between 0mCA and 2mCA infants relied on an unbiased set of microstates. Since no previous work studied microstates in 2-month-old infants, we implemented a data-driven approach to measure the templates similarity across groups and we considered shared (i.e. similar) microstates for further analyses.

Moreover, we took into account that brain activity might differ across vigilance states (Dereymaeker et al., 2017), given their crucial role in modulating developmental and learning mechanisms early in life (Li et al., 2017). We performed templates identification and analyses for each state separately. We then observed an increase in global explained variance from awake to REM sleep and further into NREM sleep, which is consistent with previous reports in full-term neonates in active vs quiet sleep (Khazaei et al., 2021) and suggests a global reduction in complexity of brain dynamics with progression in sleep depth. While such effects might vary across microstates, as shown in neonates (Khazaei et al., 2021, see also Khazaei et al., 2023) and adults (Brodbeck et al., 2012), evaluating the precise impact of vigilance states was beyond the scope of our study.

Besides, our study benefited from HD-EEG recordings that are suggested to improve reliability of microstates analysis (Zhang et al., 2021). Interestingly, the microstates we identified in preterms at TEA were similar to those of full-term asleep neonates reported in the previous study in similar sleep stages (Khazaei et al., 2021), suggesting that such approach is quite reproducible across different study settings, neonatal populations and recording systems, and only with a few minutes of recording (Bagdasarov et al., 2024). This opens up the perspective of implementing microstate templates by age group, available to the scientific and clinical community for future multi-site studies for example.

Finally, our study was exploratory in taking a step towards characterizing early maturation and dysmaturation of microstate dynamics reflecting the brain activity. Since our study was limited by the sample size of preterms (n=43 but with inter-individual variability in clinical characteristics) and full-terms (n=14) and uncomplete longitudinal evaluations, future studies in larger cohorts could better address relationships with neonatal factors. A crucial next step is to tackle the links between microstates and the underlying cerebral networks for understanding the specific functional relevance of different microstates in the developing infant brain.

### Conclusions

This study implemented a dedicated strategy for the quantitative evaluation of brain spontaneous activity in preterm infants using the EEG microstate approach, highlighting fundamental features that describe the maturation of the brain dynamics and show sensitivity to perinatal factors. Such an approach might offer novel potential diagnostic markers for understanding the heterogeneity of atypicalities and developmental trajectories over the early post-natal period of intense neuroplasticity, when interventions building on compensatory mechanisms could improve the neurodevelopmental outcomes (Livingston & Happe, 2017). Although beyond the scope of current study, this encourages future research to perform similar analysis in larger groups of full-term neonates in order to characterize the normative developmental trajectories required for a better identification of atypicalities in preterm infants with a wide inter-individual variability.

## Abbreviations

BOLD: blood oxygenation level dependent
(m)CA: (months of) corrected age
(HD-)EEG: (high-density-) electroencephalography
(w)GA: (weeks of) gestational age
(f)MRI: (functional) magnetic resonance imaging
M / F: male / female
MS: microstate
(w)PMA: (weeks of) post-menstrual age
(N)REM: (non) rapid eye movement
TEA: term equivalent age

## Acknowledgments

This work was supported by the French National Agency for Research (ANR grant ANR-20-CE17-0014-03-PremaLocom), the Fondation Médisite (under the aegis of the Fondation de France, grant FdF-18-00092867), the IdEx Université de Paris (ANR-18-IDEX-0001), the French government as part of the France 2030 programme (grant ANR-23-IAIIU-0010, IHU Robert-Debré du Cerveau de l’Enfant), as well as UK Research and Innovation (UKRI) under the UK government’s Horizon Europe funding guarantee scheme of MSCA (Grant EP/X021947/1 to P.A.). We would like to thank Guillaume Dumas and the SoNeTAA platform team (funded by Fondation de France), as well as clinical teams at Robert-Debré Hospital (in particular Nathalie Medrano, Séverine Baron, Mireille Toquer and Magali Riche) and at NeuroSpin/UNIACT (in particular Gaëlle Mediouni and Bernadette Martins) whose collaboration made it possible to conduct this study in a clinical setting. We are also grateful for helpful discussions with Catherine Chiron, Fabrice Wallois and Nadège Roche-Labarbe. Last but not least, we would like to sincerely thank all the infants and parents who participated in this study.

## Supplementary Information

### 1. Participants information

**SI Table 1.** Detailed neonatal characteristics and clinical risk factors for each of the preterm infants. These include groups of GA at birth (GA1/GA2/GA3), sex (males/females), binarized risk of birth weight indicating small for gestational age (yes/no), binarized MRI Kidokoro score (mild/normal), as well as information regarding five categories of non-neurological complications at NICU. These complications were considered for chronic lung disease (yes/no), use of invasive mechanical ventilation for more than 1 day (yes/no), necrotizing enterocolitis (yes/no), parenteral nutrition for more than 3 weeks (yes/no) and sepsis (yes/no).

| Subjects | GA at birth | GA category | Sex | Small weight for GA | MRI Kidokoro score | Chronic lung disease | Mechanical ventilation>1day | Necrotizing enterocolitis | Parenteral nutrition>3weeks | Sepsis |
| --- | --- | --- | --- | --- | --- | --- | --- | --- | --- | --- |
| 1 | 26.6 | 2 | M | No | Normal | No | No | No | No | Yes |
| 2 | 27.0 | 2 | F | No | Normal | No | Yes | Yes | No | No |
| 3 | 27.3 | 2 | F | No | Normal | No | No | No | No | Yes |
| 4 | 27.9 | 2 | M | No | Mild | No | No | No | No | No |
| 5 | 25.7 | 1 | M | No | Normal | No | No | No | No | Yes |
| 6 | 25.7 | 1 | M | No | Normal | No | No | No | No | Yes |
| 7 | 25.1 | 1 | F | Yes | Normal | Yes | Yes | No | Yes | Yes |
| 8 | 26.4 | 2 | M | No | Normal | No | No | No | Yes | Yes |
| 9 | 26.4 | 2 | F | No | Mild | Yes | No | No | Yes | Yes |
| 10 | 26.6 | 2 | M | No | Normal | No | No | No | No | Yes |
| 11 | 26.6 | 2 | M | No | Normal | No | No | No | No | Yes |
| 12 | 29.9 | 3 | M | Yes | Normal | No | Yes | Yes | Yes | Yes |
| 13 | 29.7 | 3 | F | No | Normal | No | No | No | No | Yes |
| 14 | 25.9 | 1 | F | No | Normal | No | No | No | No | Yes |
| 15 | 25.9 | 1 | M | No | Normal | Yes | No | No | Yes | Yes |
| 16 | 25.9 | 1 | F | No | Mild | No | No | No | No | Yes |
| 17 | 27.4 | 2 | M | No | Mild | Yes | No | No | No | Yes |
| 18 | 30.9 | 3 | M | No | Normal | No | No | No | No | No |
| 19 | 26.0 | 1 | M | No | Normal | Yes | No | No | Yes | Yes |
| 20 | 25.1 | 1 | F | No | Normal | Yes | Yes | No | No | Yes |
| 21 | 27.9 | 2 | M | No | Normal | Yes | No | No | No | No |
| 22 | 28.1 | 3 | F | Yes | Normal | Yes | Yes | Yes | Yes | Yes |
| 23 | 27.7 | 2 | M | No | Normal | No | No | No | No | No |
| 24 | 28.9 | 3 | F | No | Normal | Yes | Yes | No | Yes | Yes |
| 25 | 29.7 | 3 | M | No | Normal | Yes | Yes | No | Yes | Yes |
| 26 | 24.3 | 1 | F | No | Normal | Yes | Yes | No | Yes | Yes |
| 27 | 26.7 | 2 | F | No | Normal | Yes | No | No | No | No |
| 28 | 25.0 | 1 | F | No | Normal | Yes | Yes | Yes | Yes | Yes |
| 29 | 27.1 | 2 | M | No | Normal | Yes | Yes | No | Yes | No |
| 30 | 28.6 | 3 | M | Yes | Normal | Yes | No | No | No | Yes |
| 31 | 29.4 | 2 | F | No | Normal | Yes | No | No | Yes | Yes |
| 32 | 27.6 | 2 | M | Yes | Normal | Yes | No | No | No | Yes |
| 33 | 26.3 | 2 | F | No | Normal | Yes | Yes | No | Yes | Yes |
| 34 | 26.7 | 2 | M | No | Normal | Yes | No | No | No | Yes |
| 35 | 28.3 | 3 | M | No | Normal | Yes | Yes | No | Yes | Yes |
| 36 | 24.6 | 1 | M | Yes | Normal | Yes | Yes | No | Yes | Yes |
| 37 | 24.1 | 1 | M | No | Mild | Yes | Yes | Yes | Yes | Yes |
| 38 | 30.6 | 3 | M | Yes | Mild | No | No | No | No | No |
| 39 | 26.3 | 2 | F | No | Normal | Yes | Yes | Yes | Yes | Yes |
| 40 | 27.1 | 2 | F | No | Normal | Yes | Yes | No | Yes | No |
| 41 | 28.1 | 3 | F | No | Mild | No | No | No | No | No |
| 42 | 25.1 | 1 | F | No | Normal | Yes | Yes | Yes | Yes | Yes |
| 43 | 30.9 | 3 | F | Yes | Normal | No | No | No | No | No |

### 2. Comparing microstates between preterm and full-term infants: Impact of prematurity on MS coverage and occurrence and global explained variance

**SI Table 2.**
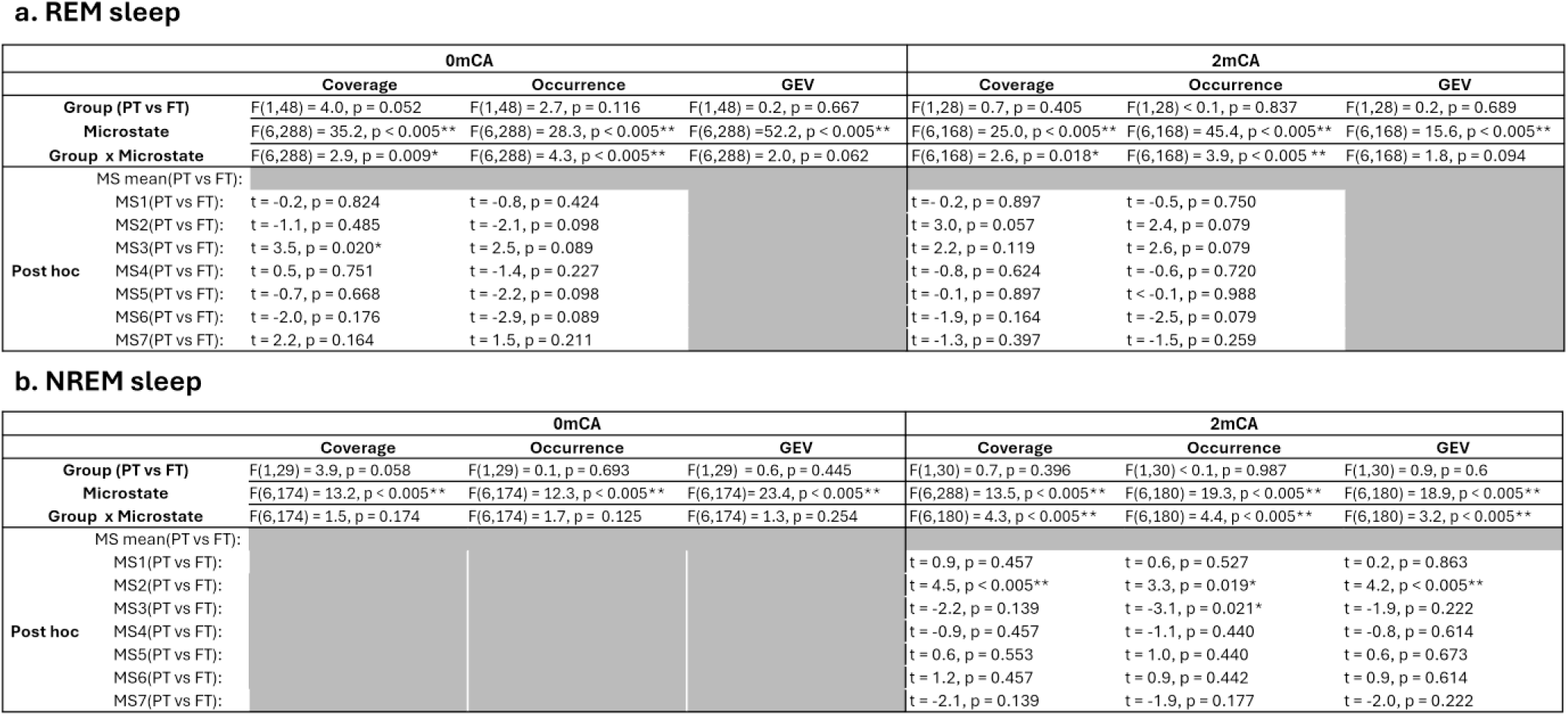
Comparison of microstates metrics (coverage, occurrence and global explained variance-GEV) between Preterm (PT) and Full-Term (FT) infants for different sleep states. Statistical tests were performed for n=41/n=22 preterm vs n=9/n=8 full-term at 0mCA/2mCA in REM sleep as well as for n=21/n=23 preterm vs n=10/n=9 full-term infants at 0mCA/2mCA in NREM sleep. P values were corrected for multiple comparisons with FDR approach for each set of post hoc tests, and significant statistical tests are indicated with asterisks (* p<0.05, ** p<0.005).

**SI Figure 1.**
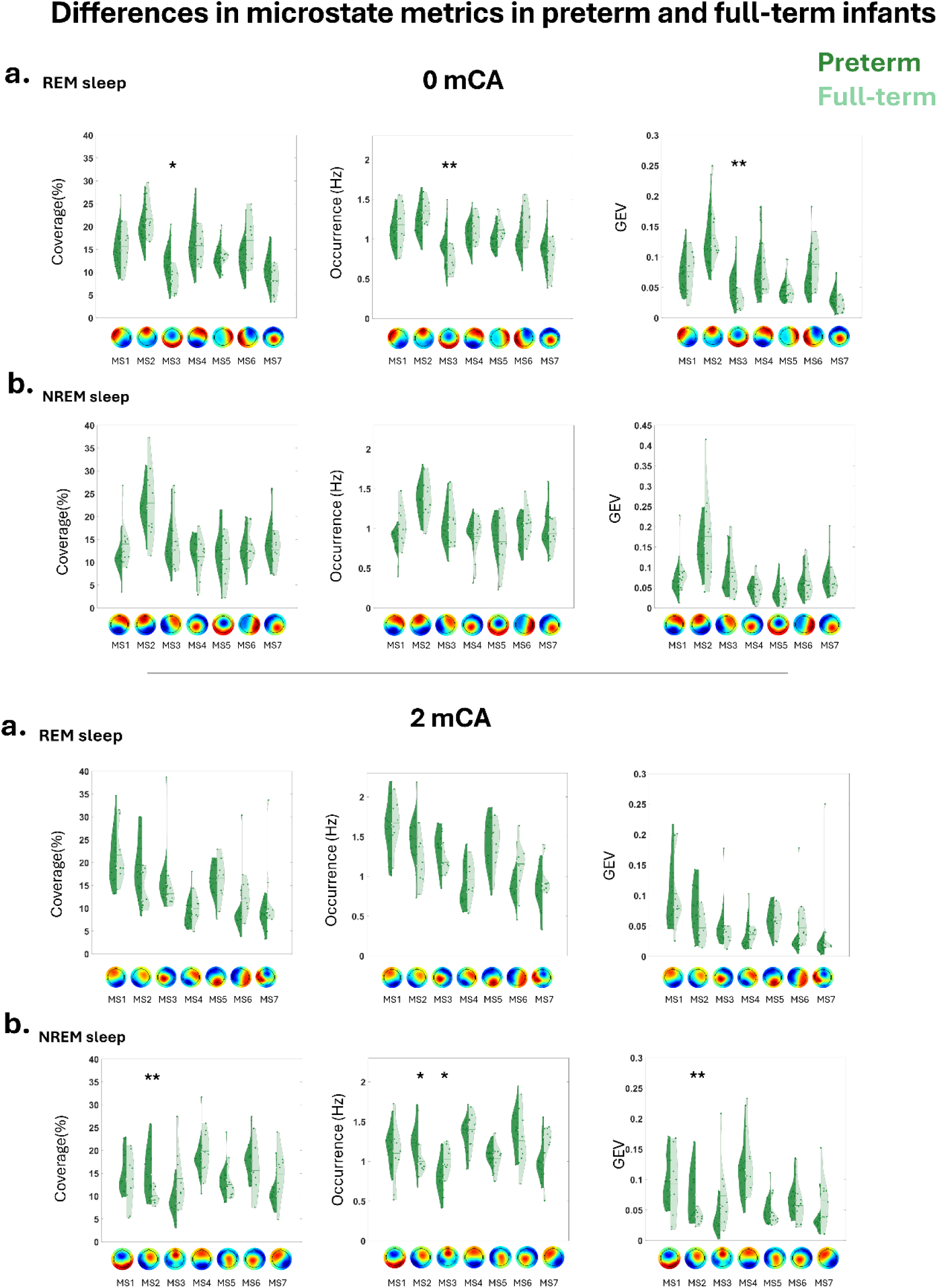
Changes in microstate metrics (coverage, occurrence and global explained variance-GEV) between preterms (dark green) and full-terms (light green) in **a.** REM sleep; **b.** NREM sleep. Asterisks represent significant differences between preterm and full-term infants (** p<0.005, *p<0.05).

### 3. Comparing microstates between 0mCA and 2mCA: Impact of maturation on MS dynamics in preterms

#### 3.1. Maturation of individual microstate metrics

**SI Table 3.**
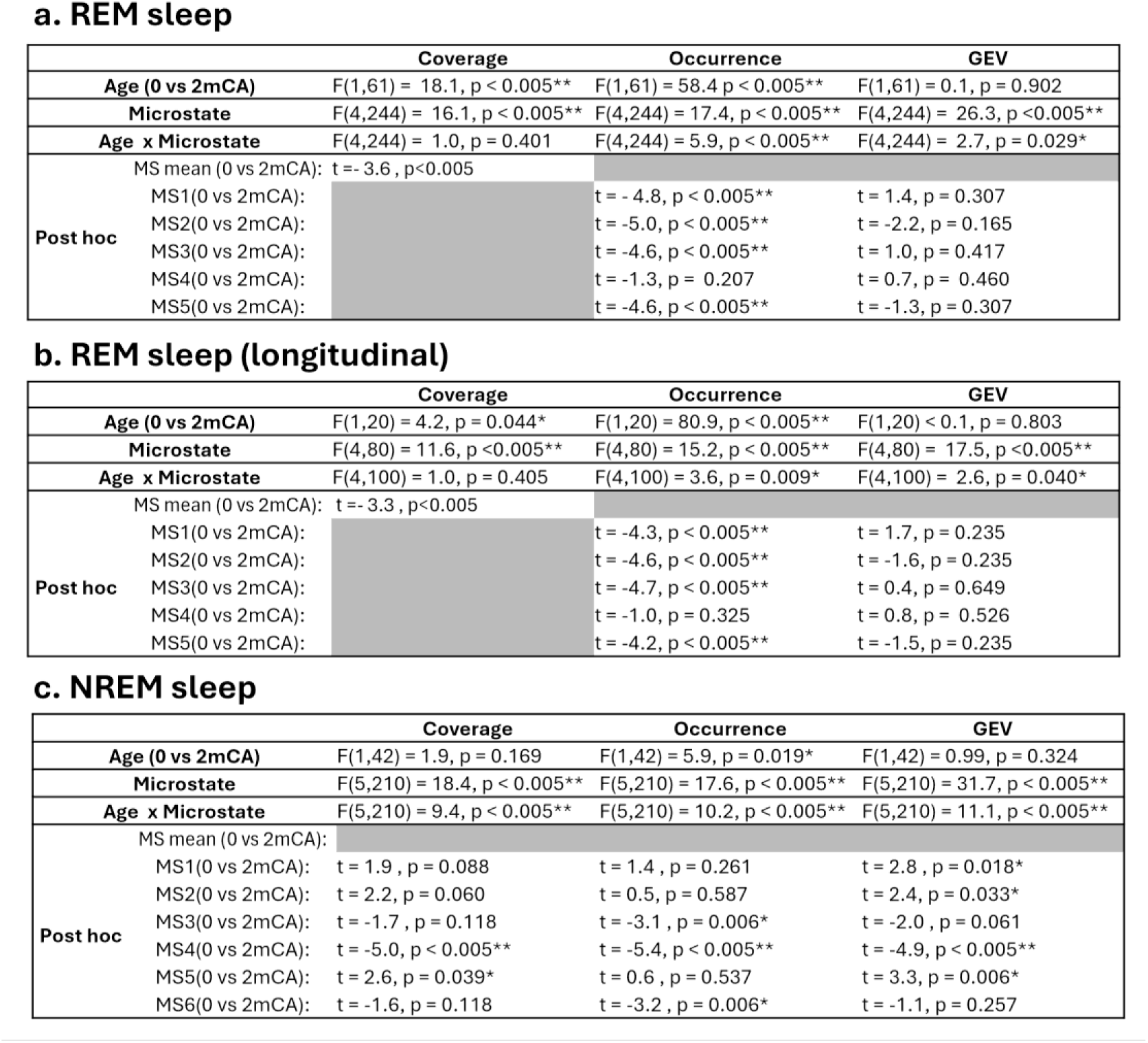
Maturation of microstate metrics (coverage, occurrence and global explained variance-GEV) between 0mCA and 2mCA during sleep states. . Statistical comparisons were performed for n=41 at 0mCA vs n=22 at 2mCA preterm infants during REM and for n=21 at 0mCA vs n=23 at 2mCA preterm infants during NREM sleep. The statistical tests in a. and c. were made cross-sectionally and in b. for n=21 infants longitudinally. P values were corrected for multiple comparisons with FDR approach for each set of post hoc tests, and significant statistical tests are indicated with asterisk (* p<0.05, ** p<0.005).

**SI Figure 2.**
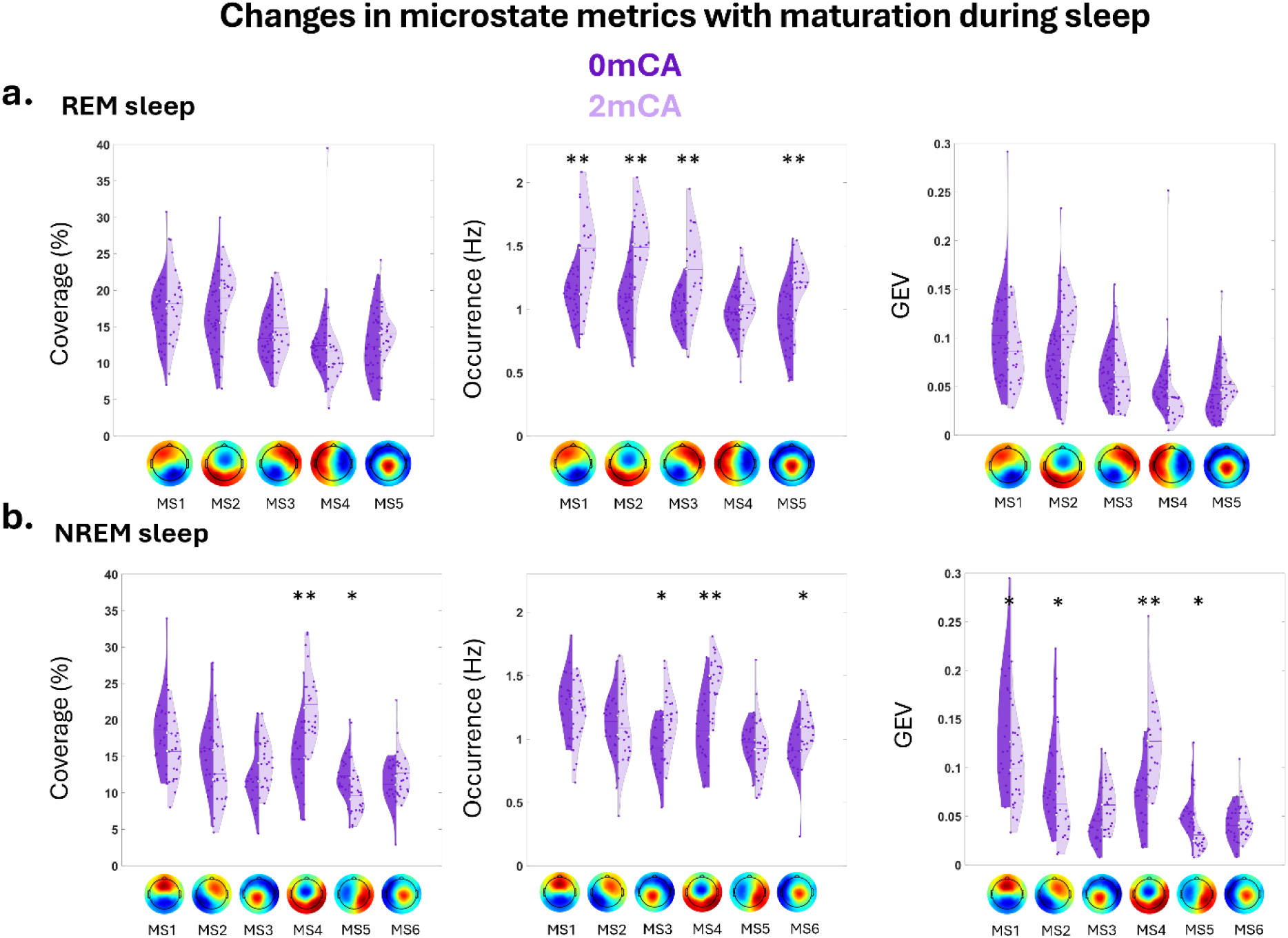
Evolution in microstate metrics (coverage, occurrence and global explained variance-GEV) between 0mCA (dark purple) and 2mCA (light purple) in preterms, during different sleep states for the **a.** five shared microstates in REM sleep; **b.** six shared microstates in NREM sleep. Asterisks represent significant differences between 0mCA and 2mCA infants (** p<0.005, *p<0.05).

#### 3.2. Maturation of microstates transitions

Besides metrics describing individual microstates, we also explored the transitions between microstates, and quantified the transition probabilities of microstates, indicating how frequently microstates of a certain class are followed by microstates of other classes (transition probability of X⇒Y: number of times X is followed by Y divided by the number of total transitions). Since transition probabilities between microstates also depend on the base occurrence rate of each microstate, we divided (i.e., normalized) the transition probabilities by the occurrence rate of the ‘destination’ microstates (i.e., the microstate that the transition goes into). To assess whether each microstate has favorable microstates to transition into at each age, we compared the normalized transition probability between a pair of microstates (i.e., an initial microstate and a destination microstate) and the alternative transitions from the initial microstate using paired t-tests: the transition was considered favorable if higher than at least half of the alternative transitions. To have comparable results between the two ages and evaluate whether such transitioning patterns change with development, we focused on the shared microstates between the two age groups for each vigilance state. We corrected t-test statistics for multiple comparisons using the False Discovery Rate (FDR) approach.

**SI Figure 3.**
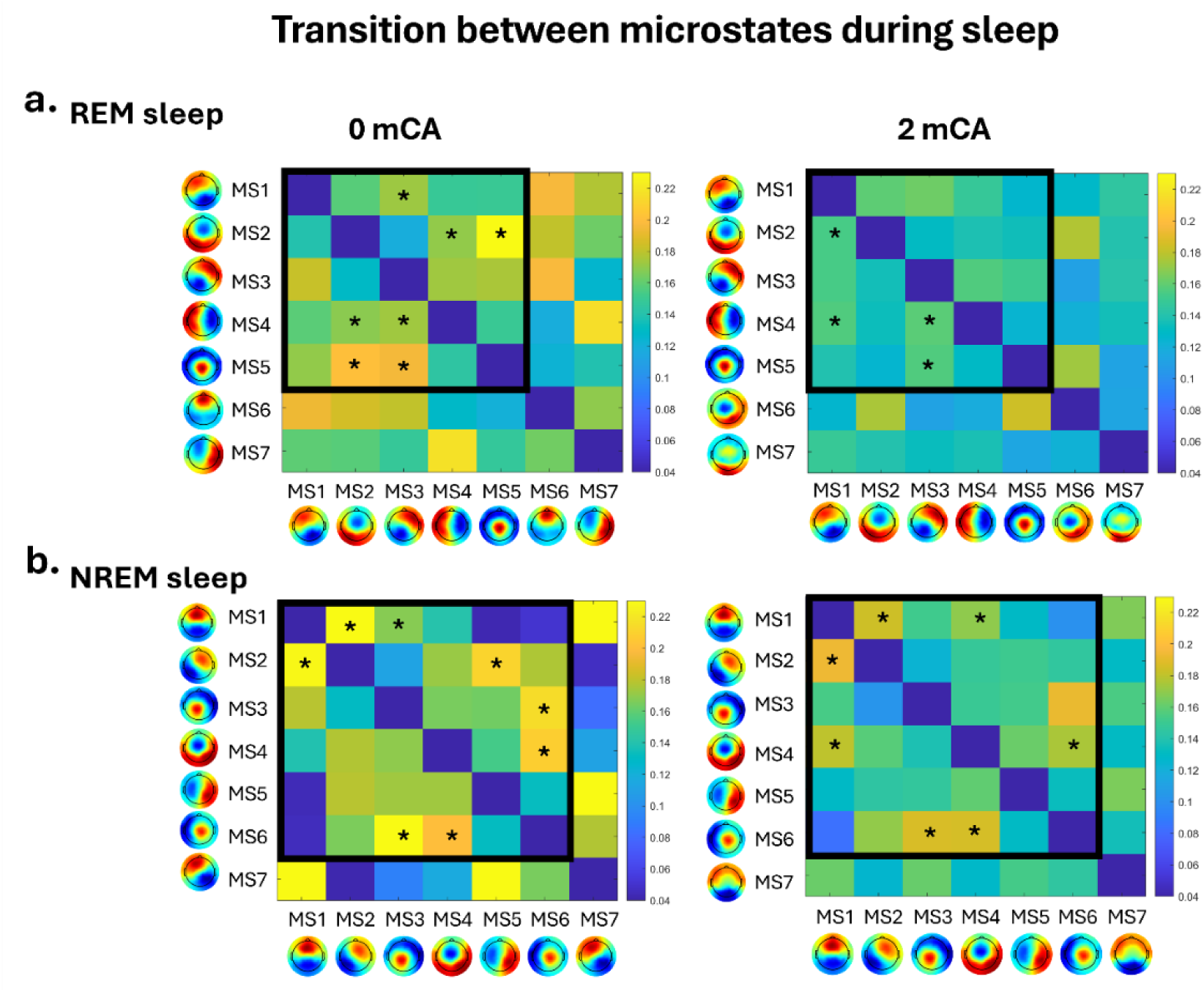
Normalized transition probabilities between different microstates at different sleep states in preterms at 0 and 2mCA (left/right column): a. REM sleep b. NREM sleep. Asterisks represent significant favorable transitions from MS XèMSY (MS X represented in rows and MS Y represented in columns). The dark squares highlight the shared microstates between 0 and 2mCA for each vigilance state.

Comparing the normalized transition probability of pairs of microstates indicated that several microstates had significant favorable transitions, whose number varied with age and sleep state (from 4 to 8). These transitions were in part stable between 0 and 2mCA for each sleep state. For REM sleep, two of the transitions remained stable between the two ages (out of the 7/4 favorable transitions identified at 0/2mCA) (SI Figure 3.a), while for NREM sleep, five transitions remained stable (out of the 8/7 favorable transitions identified at 0/2mCA) (SI Figure 3.b).

These results indicate that at both ages, the transitions between the microstates had a non-random structure, with most microstates tending to transition into their ‘favorite’ microstates (one or more than one). The pattern of such transitions evolved with development between 0 and 2mCA which could suggest a reorganization of the brain functional dynamics, showing an evolution in the continuous shifting between transient network activities. Note that although the presence of non-random structure in the transition between microstates is widely accepted, test-retest reliability assessments of ‘favorite’ transitions have been described as poor (Antonova et al., 2022; Kleinert et al., 2023; but see Liu et l., 2020; Jun et al., 2024), limiting their interpretation for capturing interindividual differences. Future work is needed to better characterize the dynamic transitions between states, perhaps by studying the brain states in finer details beyond sleep/wake states (e.g., during or following different sensory stimulation contexts). Moreover, with the number of possible transitions between the states, studying these aspects of microstate dynamics would require a larger sample of infants for reliable statistical testing.

### 4. Complementary analysis relating microstates to clinical factors

#### 4.1. Relating microstate duration at 0mCA to GA at birth across all preterm and full-term infants in REM sleep

In a more focused analysis on the impact of GA at birth on microstates duration, we related the mean duration of microstates (averaged across all 7 microstates) at 0mCA to the group of GA at birth (GA1/GA2/GA3/Full-term) using an ANOVA and follow up t-tests. This analysis confirmed an impact of GA group on microstate duration (F(3,46) = 3.1, p= 0.037)) and posthoc tests indicated that full-term neonates have shorter microstate duration than all other preterm GA groups (p<0.05 for GA1 and GA3 compared to Full-terms, and p<0.1 for GA2 compared to full-terms).

**SI Figure 4.**
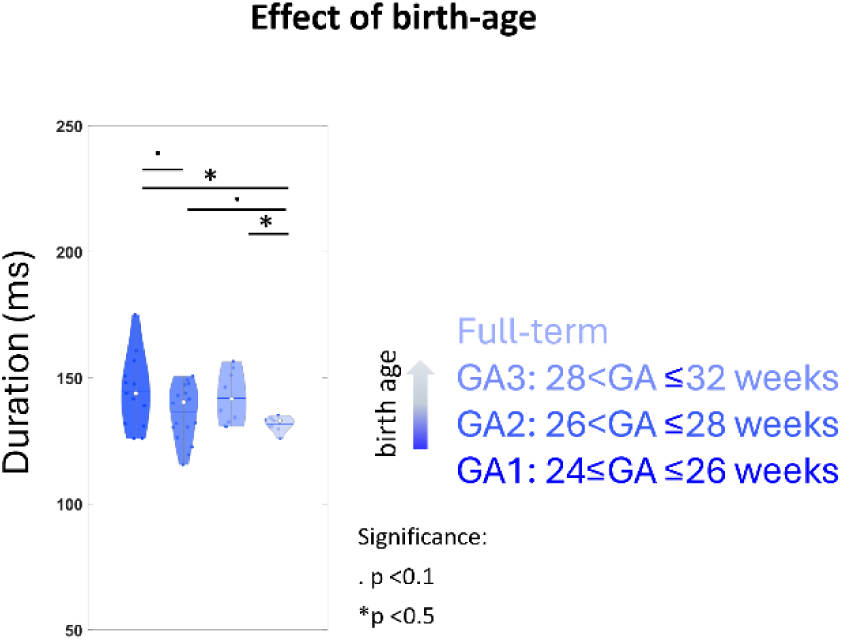
Impact of Groups of GA at birth on average MS duration in REM sleep. Asterisk and dots represent significant (p<0.05) and trend (p<0.1) differences between GA groups: longer duration in infants with the lowest GA at birth.

#### 4.2. Relating microstate duration at REM sleep and clinical factors in preterms

We performed two complementary ANOVAs similar to section 2.6.3, in order to explore the relationships between microstates duration and the prematurity risk factors. Instead of considering a binary morbidity score describing non-neurological complications during the NICU period, we considered a continuous ratio variable [0:0.2:1] summarizing how many of the five factors (chronic lung disease, need for invasive mechanical ventilation lasting strictly more than 1 day, necrotizing enterocolitis, need for parenteral nutrition longer than 3 weeks, and experiencing sepsis), were present in each infant. Due to the limited number of infants, we had to consider the average duration of microstates as dependent variable (thus the microstate factor was not included in the analysis). The remaining dependent variables (GA group, MRI score, Sex, Small weight for GA) were similar to the previous analysis in 2.6.3. The following results showed no effect, in line with the results of the analysis with the binary factor.

**SI Table 4.**
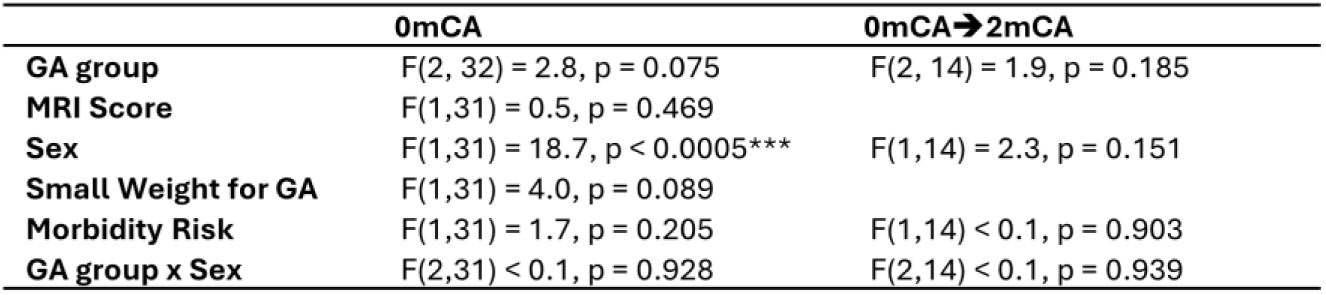
Impact of perinatal factors on microstates at 0mCA and their maturation between 0mCA and 2mCA. The relationships between microstate duration at 0mCA (n=41, left column) or the longitudinal changes in microstate duration between 0mCA and 2mCA (n=21, right column) and different prematurity risk factors were evaluated for REM sleep activity. These factors included group of GA at birth (GA1/GA2/GA3); sex; binarized risk of birth weight indicating small for gestational age; MRI Kidokoro score and continuous neonatal morbidity score summarizing non-neurological complications. At 2mCA, only main effects and interactions with adequate data points were considered. P values were corrected for multiple comparisons with FDR approach for each set of post hoc tests, and significant statistical tests are indicated with asterisk (*** p<0.0005).

|  | 0mCA | 0mCA→2mCA |
| --- | --- | --- |
| **GA group** | F(2, 32) = 2.8, p = 0.075 | F(2, 14) = 1.9, p = 0.185 |
| **MRI Score** | F(1,31) = 0.5, p = 0.469 |  |
| **Sex** | F(1,31) = 18.7, p < 0.0005*** | F(1,14) = 2.3, p = 0.151 |
| **Small Weight for GA** | F(1,31) = 4.0, p = 0.089 |  |
| **Morbidity Risk** | F(1,31) = 1.7, p = 0.205 | F(1,14) < 0.1, p = 0.903 |
| **GA group x Sex** | F(2,31) < 0.1, p = 0.928 | F(2,14) < 0.1, p = 0.939 |

### 5. Comparing microstates between 0mCA and 2mCA: Impact of maturation on MS dynamics in preterms during wakefulness

**SI Figure 5.**
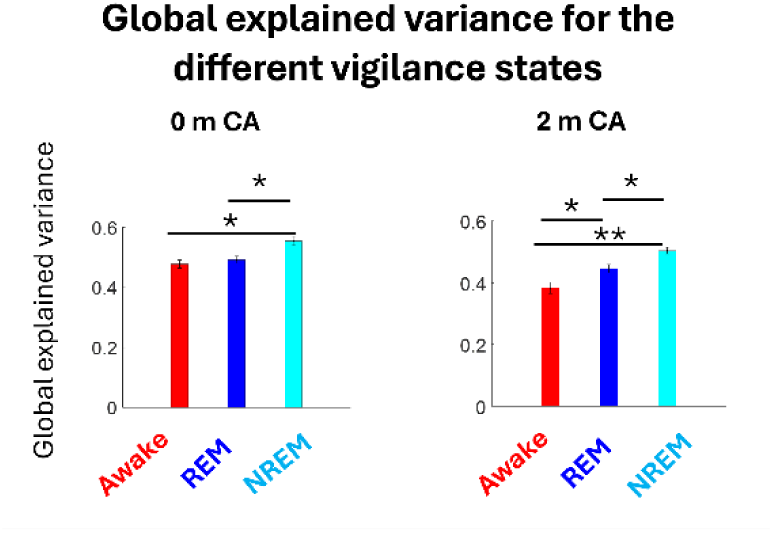
Global explained variance for the different vigilance states with 7 microstate classes. The global explained variance increased with progression in sleep depth at both ages. Significant comparisons are highlighted with asterisks (** p<0.005, * p<0.05).

**SI Table 5.**
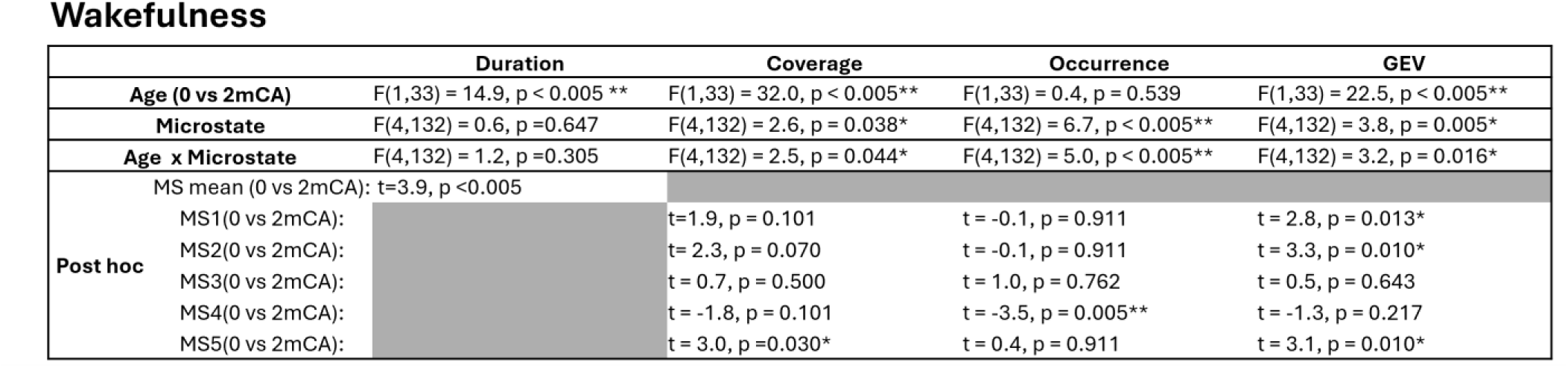
Maturation of microstate metrics (duration, global explained variance-GEV, coverage and occurrence) between 0mCA and 2mCA during wakefulness. . Statistical comparisons were performed between 0mCA (n=17) and 2mCA (n=18): P values were corrected for multiple comparisons with FDR approach for each set of post hoc tests, and significant statistical tests are indicated with asterisk (* p<0.05, ** p<0.005).

**SI Figure 6.**
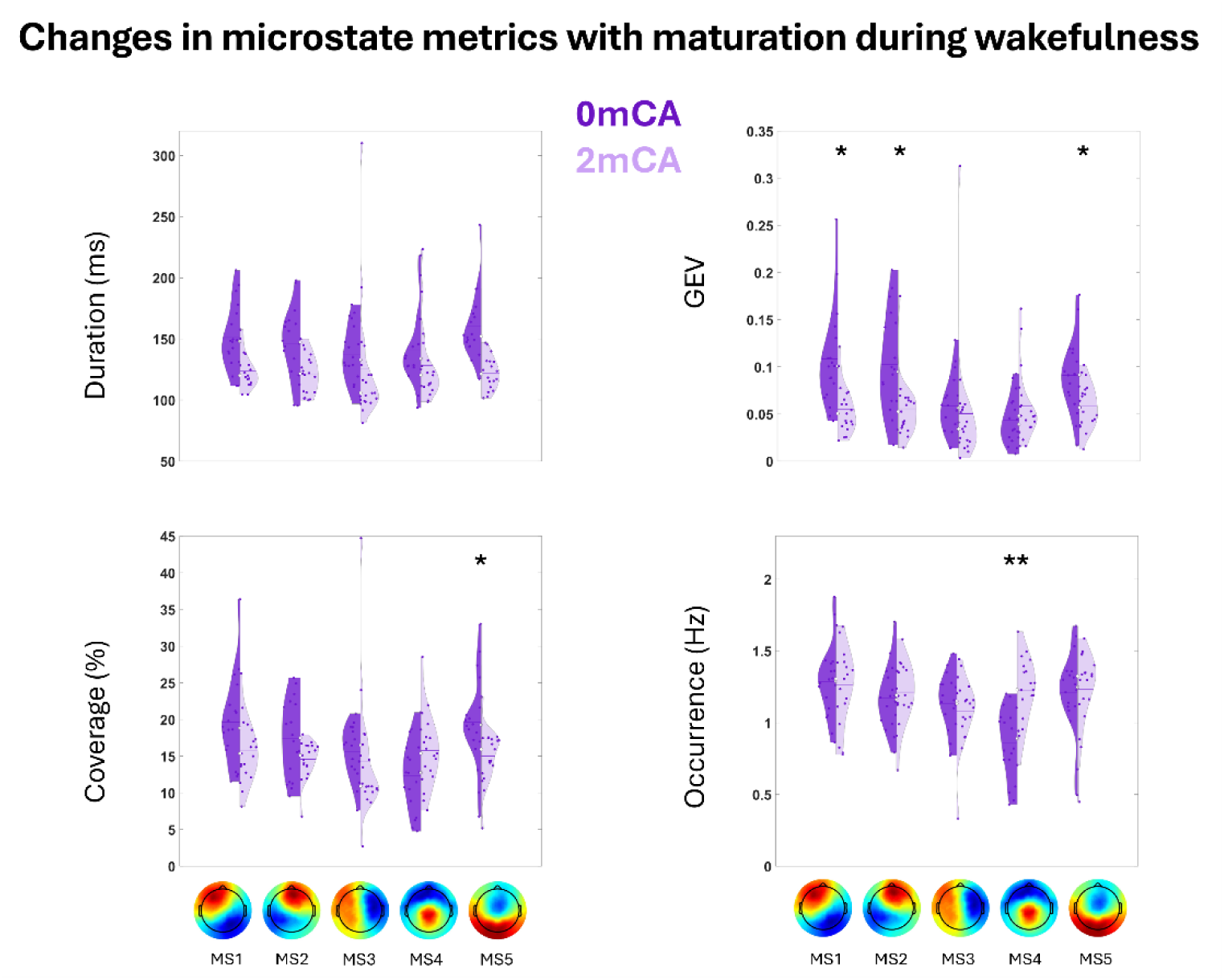
Evolution in microstate metrics (duration, global explained variance-GEV, coverage and occurrence) between 0mCA (dark purple) and 2mCA (light purple) in preterms, during wakefulness. Asterisks represent significant differences between 0mCA and 2mCA infants (** p<0.005, * p<0.05).

**SI Figure 7.**
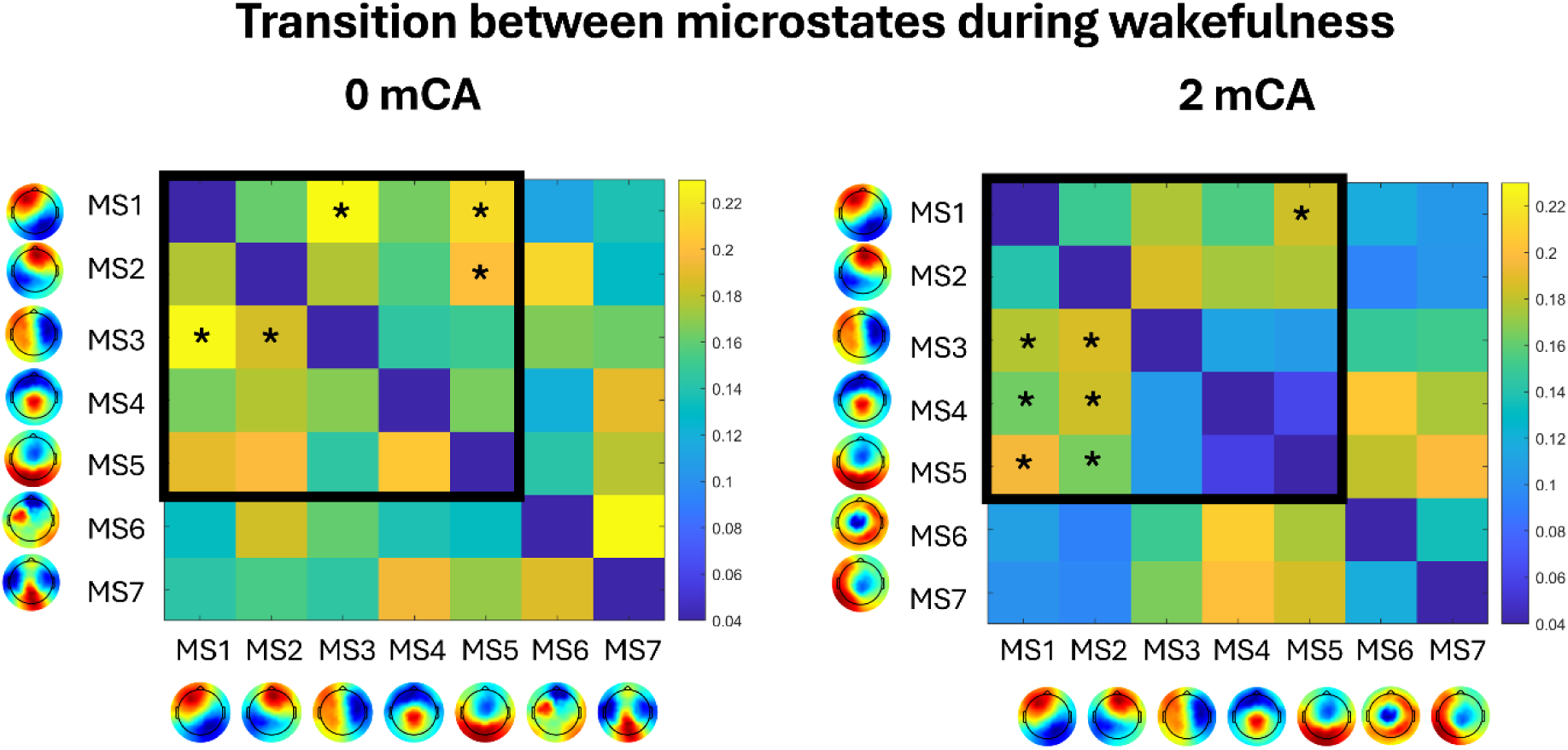
Normalized transition probabilities between different microstates during wakefulness in preterms at 0 and 2mCA (left/right column). Asterisks represent significant favorable transitions. The dark squares highlight the shared microstates between 0 and 2mCA wakefulness.

### 6. Summary of different microstate templates

**SI Figure 8.**
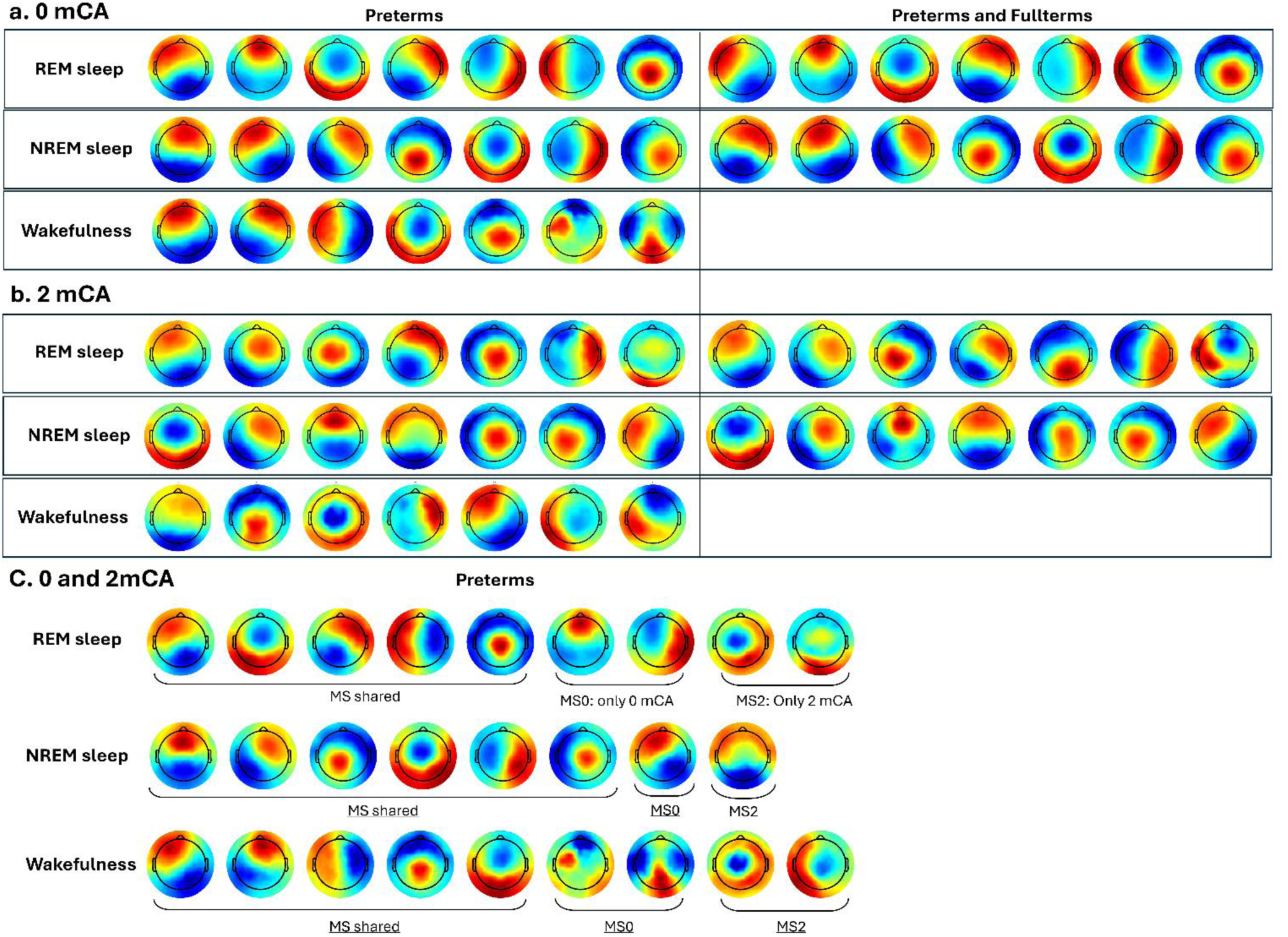
Group-level template microstates for **a.** 0mCA and **b.** 2mCA age groups across different vigilance states. Template microstates obtained considering only the preterm group are presented in the left panel, and those obtained by considering preterm and full-term infants are presented in the right panel. **c.** Group-level template microstates obtained in preterms when considering both 0mCA and 2mCA groups following the procedure described in Figure 2. These microstates involve shared patterns (“MS shared”) between both age groups as well as non-shared ones (i.e. non-similar) for each age group (0mCA/2mCA → MS0/MS2).

### 7. Ranges of different microstate metrics

**SI Table 6.** Descriptive summary of microstate metrics for preterms and fullterms during REM and NREM sleep at 0 and 2mCA. Mean, ±standard deviation and [range] are indicated.

a. REM – 0mCA
|  | Duration(msec) |  |  |  | Coverage(%) |  |  |  | Occurrence (Hz) |  |  |  | GEV (ratio) |  |  |  |
| --- | --- | --- | --- | --- | --- | --- | --- | --- | --- | --- | --- | --- | --- | --- | --- | --- |
|  | preterms |  | fullterms |  | preterms |  | fullterms |  | preterms |  | fullterms |  | preterms |  | fullterms |  |
| | mean±std | [min max] | mean±std | [min max] | mean±std | [min max] | mean±std | [min max] | mean±std | [min max] | mean±std | [min max] | mean±std | [min max] | mean±std | [min max] |
| MS1 | 137±19 | [104 194] | 130±11 | [109 144] | 15±3 | [8 27] | 15±4 | [8 21] | 1.10±0.19 | [0.75 1.49] | 1.18±0.26 | [0.75 1.55] | 0.07±0.03 | [0.031 0.145] | 0.07±0.03 | [0.02 0.12] |
| MS2 | 160±19 | [122 221] | 158±15 | [141 166] | 20±3 | [12 29] | 21±4 | [16 29] | 1.24±0.19 | [0.87 1.65] | 1.35±0.14 | [1.17 1.59] | 0.12±0.04 | [0.063 0.235] | 0.13±0.04 | [0.07 0.25] |
| MS3 | 128±13 | [101 164] | 108±12 | [92 127] | 11±3 | [4 20] | 8±2 | [4 12] | 0.89±0.23 | [0.41 1.49] | 0.73±0.16 | [0.52 0.94] | 0.05±0.03 | [0.008 0.133] | 0.02±0.01 | [0.01 0.05] |
| MS4 | 154±27 | [111 221] | 137±14 | [121 158] | 16±4 | [7 28] | 15±3 | [11 21] | 1.05±0.16 | [0.69 1.46] | 1.13±0.16 | [0.90 1.38] | 0.08±0.04 | [0.026 0.183] | 0.06±0.02 | [0.04 0.12] |
| MS5 | 130±15 | [100 156] | 123±11 | [104 148] | 13±2 | [8 19] | 13±2 | [9 20] | 1.00±0.13 | [0.78 1.29] | 1.11±0.14 | [0.89 1.37] | 0.04±0.01 | [0.024 0.096] | 0.04±0.02 | [0.02 0.09] |
| MS6 | 133±21 | [92 194] | 136±19 | [113 160] | 13±3 | [6 24] | 16±5 | [10 25] | 0.97±0.18 | [0.60 1.42] | 1.21±0.23 | [0.88 1.56] | 0.06±0.03 | [0.026 0.183] | 0.08±0.03 | [0.04 0.14] |
| MS7 | 120±15 | [87 155] | 105±11 | [86 118] | 11±3 | [3 18] | 8±3 | [3 12] | 0.88±0.22 | [0.38 1.48] | 0.75±0.22 | [0.40 1.03] | 0.03±0.01 | [0.005 0.074] | 0.02±0.01 | [0.01 0.03] |

b. NREM – 0mCA
|  | Duration(msec) |  |  |  | Coverage(%) |  |  |  | Occurrence (Hz) |  |  |  | GEV (ratio) |  |  |  |
| --- | --- | --- | --- | --- | --- | --- | --- | --- | --- | --- | --- | --- | --- | --- | --- | --- |
|  | preterms |  | fullterms |  | preterms |  | fullterms |  | preterms |  | fullterms |  | preterms |  | fullterms |  |
| | mean±std | [min max] | mean±std | [min max] | mean±std | [min max] | mean±std | [min max] | mean±std | [min max] | mean±std | [min max] | mean±std | [min max] | mean±std | [min max] |
| MS1 | 131±24 | [90 181] | 141±251 | [116 179] | 14±5 | [5 24] | 17±6 | [10 28] | 1.06±0.25 | [0.56 1.42] | 1.20±0.24 | [0.83 1.63] | 0.09±0.05 | [0.02 0.22] | 0.12±0.05 | [0.06 0.23] |
| MS2 | 153±27 | [100 227] | 156±41 | [89 238] | 19±5 | [8 28] | 22±9 | [6 40] | 1.28±0.24 | [0.84 1.75] | 1.36±0.30 | [0.77 1.78] | 0.14±0.05 | [0.04 0.24] | 0.17±0.11 | [0.02 0.40] |
| MS3 | 122±21 | [86 166] | 118±18 | [94 148] | 11±3 | [3 20] | 12±4 | [7 18] | 0.94±0.18 | [0.36 1.23] | 1.05±0.21 | [0.74 1.39] | 0.05±0.02 | [0.02 0.11] | 0.06±0.03 | [0.02 0.11] |
| MS4 | 130±26 | [96 191] | 116±18 | [90 137] | 11±4 | [2 21] | 9±4 | [3 16] | 0.89±0.24 | [0.25 1.24] | 0.79±0.28 | [0.33 1.19] | 0.04±0.02 | [0.00 0.11] | 0.03±0.02 | [0.00 0.06] |
| MS5 | 149±20 | [110 183] | 134±24 | [80 168] | 17±5 | [7 26] | 13±4 | [2 18] | 1.15±0.27 | [0.67 1.71] | 0.97±0.28 | [0.22 1.25] | 0.08±0.04 | [0.02 0.18] | 0.05±0.02 | [0.00 0.09] |
| MS6 | 125±20 | [87 171] | 120±12 | [102 145] | 11±3 | [5 19] | 11±2 | [6 15] | 0.94±0.18 | [0.65 1.20] | 0.97±0.16 | [0.61 1.21] | 0.05±0.02 | [0.01 0.12] | 0.05±0.02 | [0.02 0.11] |
| MS7 | 132±19 | [88 164] | 125±13 | [102 150] | 12±3 | [4 17] | 12±3 | [6 17] | 0.97±0.18 | [0.39 1.25] | 0.98±0.19 | [0.63 1.19] | 0.05±0.02 | [0.01 0.10] | 0.05±0.02 | [0.01 0.10] |

c. REM – 2mCA
|  | Duration(msec) |  |  |  | Coverage(%) |  |  |  | Occurrence (Hz) |  |  |  | GEV (ratio) |  |  |  |
| --- | --- | --- | --- | --- | --- | --- | --- | --- | --- | --- | --- | --- | --- | --- | --- | --- |
|  | preterms |  | fullterms |  | preterms |  | fullterms |  | preterms |  | fullterms |  | preterms |  | fullterms |  |
| | mean±std | [min max] | mean±std | [min max] | mean±std | [min max] | mean±std | [min max] | mean±std | [min max] | mean±std | [min max] | mean±std | [min max] | mean±std | [min max] |
| MS1 | 132±22 | [95 199] | 128±19 | [109 166] | 21±5 | [13 34] | 21±6 | [14 31] | 1.60±0.35 | [1.01 2.19] | 1.66±0.30 | [1.10 2.09] | 0.10±0.05 | [0.04 0.21] | 0.10±0.06 | [0.02 0.20] |
| MS2 | 131±27 | [93 199] | 115±13 | [101 137] | 19±6 | [8 30] | 13±4 | [9 19] | 1.47±0.29 | [0.73 2.18] | 1.18±0.29 | [0.77 1.67] | 0.08±0.03 | [0.01 0.14] | 0.04±0.02 | [0.01 0.08] |
| MS3 | 120±31 | [86 246] | 111±11 | [99 131] | 16±5 | [10 38] | 13±2 | [10 17] | 1.35±0.22 | [0.85 1.66] | 1.17±0.13 | [1.01 1.42] | 0.05±0.03 | [0.02 0.17] | 0.03±0.01 | [0.01 0.05] |
| MS4 | 100±16 | [80 158] | 102±6 | [91 110] | 8±3 | [5 18] | 9±2 | [4 14] | 0.88±0.25 | [0.55 1.40] | 0.96±0.24 | [0.53 1.30] | 0.03±0.02 | [0.01 0.10] | 0.03±0.01 | [0.01 0.05] |
| MS5 | 117±14 | [95 150] | 117±10 | [1023 134] | 16±3 | [7 21] | 16±4 | [9 22] | 1.40±0.29 | [0.76 1.86] | 1.40±0.29 | [0.84 1.77] | 0.06±0.02 | [0.01 0.09] | 0.05±0.02 | [0.02 0.09] |
| MS6 | 98±28 | [73 212] | 105±8 | [91 118] | 9±5 | [3 30] | 12±3 | [6 17] | 0.87±0.19 | [0.45 1.43] | 1.15±0.29 | [0.63 1.63] | 0.03±0.03 | [0.00 0.17] | 0.04±0.02 | [0.01 0.08] |
| MS7 | 100±9 | [90 124] | 120±52 | [91 248] | 8±2 | [3 13] | 12±8 | [7 33] | 0.84±0.24 | [0.33 1.26] | 0.99±0.23 | [0.79 1.40] | 0.01±0.00 | [0.00 0.04] | 0.04±0.08 | [0.01 0.25] |

d. NREM – 2mCA
|  | Duration(msec) |  |  |  | Coverage(%) |  |  |  | Occurrence (Hz) |  |  |  | GEV (ratio) |  |  |  |
| --- | --- | --- | --- | --- | --- | --- | --- | --- | --- | --- | --- | --- | --- | --- | --- | --- |
|  | preterms |  | fullterms |  | preterms |  | fullterms |  | preterms |  | fullterms |  | preterms |  | fullterms |  |
| | mean±std | [min max] | mean±std | [min max] | mean±std | [min max] | mean±std | [min max] | mean±std | [min max] | mean±std | [min max] | mean±std | [min max] | mean±std | [min max] |
| MS1 | 128±23 | [99 182] | 118±20 | [90 148] | 15±4 | [9 22] | 13±5 | [5 21] | 1.19±0.22 | [0.76 1.63] | 1.10±0.38 | [0.52 1.73] | 0.09±0.03 | [0.04 0.17] | 0.09±0.05 | [0.01 0.17] |
| MS2 | 124±27 | [92 196] | 101±14 | [88 135] | 15±5 | [8 25] | 10±1 | [7 13] | 1.23±0.26 | [0.67 1.71] | 0.99±0.13 | [0.80 1.23] | 0.07±0.03 | [0.02 0.16] | 0.03±0.01 | [0.02 0.05] |
| MS3 | 109±18 | [73 157] | 129±39 | [92 218] | 9±3 | [3 15] | 13±6 | [7 27] | 0.80±0.22 | [0.41 1.21] | 1.02±0.16 | [0.75 1.26] | 0.03±0.02 | [0.00 0.08] | 0.07±0.05 | [0.01 0.21] |
| MS4 | 139±23 | [110 226] | 142±27 | [111 199] | 18±4 | [10 31] | 19±4 | [12 26] | 1.31±0.20 | [0.90 1.71] | 1.40±0.21 | [1.01 1.68] | 0.11±0.04 | [0.04 0.22] | 0.12±0.05 | [0.07 0.23] |
| MS5 | 119±20 | [89 186] | 120±18 | [96 151] | 13±3 | [8 24] | 12±3 | [9 18] | 1.10±0.11 | [0.82 1.31] | 1.03±0.17 | [0.75 1.35] | 0.04±0.01 | [0.02 0.11] | 0.04±0.02 | [0.02 0.08] |
| MS6 | 130±15 | [110 177] | 122±16 | [102 149] | 17±3 | [12 27] | 15±5 | [7 25] | 1.37±0.25 | [0.95 1.94] | 1.25±0.35 | [0.71 1.84] | 0.06±0.02 | [0.03 0.13] | 0.05±0.03 | [0.02 0.13] |
| MS7 | 104±15 | [83 132] | 118±23 | [95 167] | 10±3 | [6 19] | 14±5 | [4 24] | 1.01±0.22 | [0.66 1.55] | 1.21±0.28 | [0.51 1.43] | 0.03±0.02 | [0.01 0.09] | 0.06±0.04 | [0.01 0.15] |

**SI Table 7.** Descriptive summary of microstate metrics at 0mCA and 2mCA for the different vigilance states. Mean, ±standard deviation and [range] are indicated for each metric

**a. REM – preterms**
| REM –preterms | Duration(msec) |  |  |  | Coverage(%) |  |  |  | Occurrence (Hz) |  |  |  | GEV (ratio) |  |  |  |
| --- | --- | --- | --- | --- | --- | --- | --- | --- | --- | --- | --- | --- | --- | --- | --- | --- |
|  | 0mCA |  | 2mCA |  | 0mCA |  | 2mCA |  | 0mCA |  | 2mCA |  | 0mCA |  | 2mCA |  |
| | mean±std | [min max] | mean±std | [min max] | mean±std | [min max] | mean±std | [min max] | mean±std | [min max] | mean±std | [min max] | mean±std | [min max] | mean±std | [min max] |
| MS1 | 151±25 | [98 232] | 123±15 | [89 155] | 16±4 | [7 30] | 18±4 | [8 27] | 1.09±0.18 | [0.69 1.50] | 1.47±0.35 | [0.80 2.08] | 0.10±0.04 | [0.03 0.29] | 0.08±0.03 | [0.02 0.15] |
| MS2 | 147±25 | [95 250] | 126±17 | [93 157] | 16±4 | [6 30] | 18±4 | [6 25] | 1.08±0.23 | [0.55 1.68] | 1.48±0.33 | [0.62 2.04] | 0.08±0.04 | [0.01 0.23] | 0.10±0.04 | [0.01 0.17] |
| MS3 | 135±20 | [100 182] | 112±15 | [89 156] | 13±3 | [6 21] | 14±4 | [6 22] | 0.98±0.15 | [0.68 1.32] | 1.31±0.31 | [0.62 1.94] | 0.06±0.02 | [0.02 0.15] | 0.06±0.03 | [0.02 0.13] |
| MS4 | 125±17 | [91 174] | 108±40 | [79 277] | 12±2 | [6 20] | 11±6 | [3 39] | 0.97±0.15 | [0.62 1.33] | 1.03±0.23 | [0.42 1.48] | 0.04±0.01 | [0.01 0.11] | 0.04±0.04 | [0.00 0.2] |
| MS5 | 130±20 | [94 180] | 115±22 | [87 182] | 12±4 | [4 22] | 14±3 | [6 24] | 0.92±0.25 | [0.43 1.55] | 1.21±0.21 | [0.65 1.54] | 0.04±0.02 | [0.01 0.10] | 0.05±0.02 | [0.01 0.14] |
| MS6 | 150±19 | [105 197] | 117±27 | [90 200] | 15±3 | [7 25] | 14±5 | [7 30] | 1.04±0.20 | [0.63 1.58] | 1.22±0.27 | [0.62 1.61] | 0.09±0.03 | [0.02 0.22] | 0.06±0.04 | [0.01 0.16] |
| MS7 | 129±14 | [100 159] | 109±45 | [83 297] | 13±3 | [7 21] | 7±6 | [1 33] | 1.01±0.18 | [0.65 1.48] | 0.66±0.26 | [0.22 1.19] | 0.05±0.02 | [0.02 0.10] | 0.02±0.03 | [0.00 0.17] |

**b. NREM – preterms**
| NREM-preterms | Duration(msec) |  |  |  | Coverage(%) |  |  |  | Occurrence (Hz) |  |  |  | GEV (ratio) |  |  |  |
| --- | --- | --- | --- | --- | --- | --- | --- | --- | --- | --- | --- | --- | --- | --- | --- | --- |
|  | 0mCA |  | 2mCA |  | 0mCA |  | 2mCA |  | 0mCA |  | 2mCA |  | 0mCA |  | 2mCA |  |
| | mean±std | [min max] | mean±std | [min max] | mean±std | [min max] | mean±std | [min max] | mean±std | [min max] | mean±std | [min max] | mean±std | [min max] | mean±std | [min max] |
| MS1 | 143±27 | [113 245] | 131±18 | [101 175] | 18±5 | [11 33] | 15±4 | [8 24] | 1.29±0.23 | [0.91 1.81] | 1.18±0.24 | [0.85 1.55] | 0.14±0.05 | [0.06 0.29] | 0.098±0.04 | [0.03 0.20] |
| MS2 | 139±35 | [88 251] | 114±18 | [87 162] | 16±5 | [5 27] | 12±4 | [4 23] | 1.14±0.22 | [0.61 1.61] | 1.09±0.30 | [0.39 1.65] | 0.09±0.05 | [0.02 0.22] | 0.06±0.04 | [0.010 0.15] |
| MS3 | 127±22 | [95 181] | 122±14 | [98 161] | 12±4 | [4 20] | 14±3 | [8 20] | 0.92±0.20 | [0.46 1.22] | 1.15±0.23 | [0.69 1.61] | 0.04±0.02 | [0.01 0.12] | 0.06±0.02 | [0.03 0.11] |
| MS4 | 135±18 | [96 161] | 151±27 | [115 212] | 14±4 | [6 26] | 22±4 | [14 31] | 1.07±0.27 | [0.62 1.64] | 1.46±0.20 | [1.06 1.80] | 0.06±0.03 | [0.01 0.14] | 0.13±0.04 | [0.06 0.25] |
| MS5 | 125±20 | [83 171] | 104±12 | [84 132] | 12±3 | [5 20] | 9±3 | [5 19] | 0.97±0.15 | [0.63 1.21] | 0.92±0.25 | [0.53 1.62] | 0.05±0.02 | [0.01 0.12] | 0.03±0.02 | [0.00 0.08] |
| MS6 | 123±16 | [91 156] | 115±19 | [90 167] | 11±3 | [2 15] | 12±3 | [8 22] | 0.91±0.21 | [0.23 1.29] | 1.09±0.16 | [0.759 1.38] | 0.04±0.02 | [0.01 0.07] | 0.05±0.02 | [0.019 0.10] |
| MS7 | 128±18 | [88 154] | 138±45 | [85 317] | 15±4 | [6 20] | 13±5 | [7 35] | 1.16±0.23 | [0.71 1.59] | 0.93±0.13 | [0.74 1.17] | 0.09±0.04 | [0.02 0.16] | 0.06±0.05 | [0.02 0.27] |

**c. Wakefulness – preterms**
| AWAKE-preterms | Duration(msec) |  |  |  | Coverage(%) |  |  |  | Occurrence (Hz) |  |  |  | GEV (ratio) |  |  |  |
| --- | --- | --- | --- | --- | --- | --- | --- | --- | --- | --- | --- | --- | --- | --- | --- | --- |
|  | 0mCA |  | 2mCA |  | 0mCA |  | 2mCA |  | 0mCA |  | 2mCA |  | 0mCA |  | 2mCA |  |
| | mean±std | [min max] | mean±std | [min max] | mean±std | [min max] | mean±std | [min max] | mean±std | [min max] | mean±std | [min max] | mean±std | [min max] | mean±std | [min max] |
| MS1 | 151±27 | [111 206] | 123±14 | [104 157] | 19±6 | [11 36] | 15±4 | [9 26] | 1.28±0.25 | [0.86 1.87] | 1.26±0.27 | [0.76 1.67] | 0.10±0.05 | [0.04 0.25] | 0.05±0.02 | [0.02 0.12] |
| MS2 | 146±29 | [95 197] | 120±16 | [99 149] | 17±5 | [9 25] | 14±2 | [6 17] | 1.17±0.22 | [0.79 1.70] | 1.21±0.21 | [0.67 1.58] | 0.10±0.05 | [0.02 0.20] | 0.05±0.03 | [0.01 0.17] |
| MS3 | 137±24 | [96 177] | 125±51 | [81 310] | 15±3 | [7 20] | 14±8 | [2 44] | 1.13±0.20 | [0.77 1.48] | 1.08±0.24 | [0.33 1.44] | 0.05±0.03 | [0.01 0.12] | 0.05±0.06 | [0.00 0.31] |
| MS4 | 141±31 | [93 218] | 128±31 | [98 223] | 12±4 | [4 20] | 15±4 | [7 28] | 0.86±0.24 | [0.43 1.20] | 1.22±0.23 | [0.70 1.83] | 0.04±0.02 | [0.00 0.09] | 0.05±0.03 | [0.01 0.16] |
| MS5 | 160±28 | [117 243] | 12±13 | [101 146] | 19±6 | [6 33] | 15±3 | [5 23] | 1.21±0.29 | [0.49 1.67] | 1.23±0.27 | [0.44 1.58] | 0.09±0.04 | [0.01 0.17] | 0.05±0.02 | [0.01 0.10] |
| MS6 | 135±128 | [70 646] | 123±41 | [86 270] | 6±6 | [1 32] | 13±6 | [4 34] | 0.42±0.12 | [0.21 0.73] | 1.05±0.24 | [0.49 1.53] | 0.02±0.06 | [0.00 0.27] | 0.04±0.04 | [0.00 0.17] |
| MS7 | 132±27 | [88 180] | 113±21 | [89 169] | 9±3 | [5 17] | 11±4 | [4 23] | 0.69±0.15 | [0.37 1.12] | 0.89±0.25 | [0.45 1.40] | 0.02±0.01 | [0.00 0.05] | 0.03±0.02 | [0.00 0.08] |

## Notes

### Competing Interest Statement

The authors have declared no competing interest.

### Summary of Updates

This version of the manuscript has been revised with some methodological clarifications, as well as more explicit indications of the study limitations. To simplify the main body of the manuscript, some analyses are further moved to the supplementary iformation.

## References

Allen, M. C. (2008). Neurodevelopmental outcomes of preterm infants. Current opinion in neurology, 21(*2*), 123-128.

Ajayi-Obe, M., Saeed, N., Cowan, F. M., Rutherford, M. A., & Edwards, A. D. (2000). Reduced development of cerebral cortex in extremely preterm infants. The Lancet, 356(*9236*), 1162-1163.

Arpi, E., & Ferrari, F. (2013). Preterm birth and behaviour problems in infants and preschool-age children: A review of the recent literature. Developmental Medicine & Child Neurology, 55(*9*), 788-796.

Ball, G., Aljabar, P., Zebari, S., Tusor, N., Arichi, T., Merchant, N., … & Counsell, S. J. (2014). Rich-club organization of the newborn human brain. Proceedings of the National Academy of Sciences, 111(20), 456-7

Bagdasarov, A., Brunet, D., Michel, C. M., & Gaffrey, M. S. (2024). Microstate Analysis of continuous infant EEG: Tutorial and Reliability. Brain Topography, 1–18.

Bagdasarov, A., Roberts, K., Bréchet, L., Brunet, D., Michel, C. M., & Gaffrey, M. S. (2022). Spatiotemporal dynamics of EEG microstates in four-to eight-year-old children: Age-and sex-related effects. Developmental cognitive neuroscience, 57, 101134.

Baker, A. P., Brookes, M. J., Rezek, I. A., Smith, S. M., Behrens, T., Probert Smith, P. J., & Woolrich, M. (2014). Fast transient networks in spontaneous human brain activity. elife, 3, e01867.461.

Baranger, J., Demene, C., Frerot, A., Faure, F., Delanoë, C., Serroune, H., … & Tanter, M. (2021). Bedside functional monitoring of the dynamic brain connectivity in human neonates. Nature communications, 12(*1*), 1080.

Bhutta, A. T., Cleves, M. A., Casey, P. H., Cradock, M. M., & Anand, K. J. (2002). Cognitive and behavioral outcomes of school-aged children who were born preterm: a meta-analysis. Jama, 288(6), 728–737.

Bouyssi-Kobar, M., Brossard-Racine, M., Jacobs, M., Murnick, J., Chang, T., & Limperopoulos, C. (2018). Regional microstructural organization of the cerebral cortex is affected by preterm birth. NeuroImage: Clinical, 18, 871-880.

Brodbeck, V., Kuhn, A., von Wegner, F., Morzelewski, A., Tagliazucchi, E., Borisov, S., … & Laufs, H. (2012). EEG microstates of wakefulness and NREM sleep. Neuroimage, 62(*3*), 2129-2139.

Brown, K. L., & Gartstein, M. A. (2023). Microstate analysis in infancy. Infant Behavior and Development, 70, 101785. BSI | PSC | NHSN | CDC. 2024. Available on: https://www.cdc.gov/nhsn/psc/bsi/index.html

Delorme, A., & Makeig, S. (2004). EEGLAB: an open source toolbox for analysis of single-trial EEG dynamics including independent component analysis. Journal of neuroscience methods, 134(*1*), 9-21.

Dereymaeker, A., Pillay, K., Vervisch, J., De Vos, M., Van Huffel, S., Jansen, K., & Naulaers, G. (2017). Review of sleep-EEG in preterm and term neonates. Early human development, 113, 87-103.

Dimitrova, R., Pietsch, M., Ciarrusta, J., Fitzgibbon, S. P., Williams, L. Z., Christiaens, D., … ’ O’Muircheartaigh, J. (2021). Preterm birth alters the development of cortical microstructure and morphology at term-equivalent age. NeuroImage, 243, 118488.

Dimitrova, R., Pietsch, M., Christiaens, D., Ciarrusta, J., Wolfers, T., Batalle, D., … & O’Muircheartaigh, J. (2020). Heterogeneity in brain microstructural development following preterm birth. Cerebral Cortex, 30(*9*), 4800-4810.

Dubois, J., Dehaene-Lambertz, G., Kulikova, S., Poupon, C., Hüppi, P. S., & Hertz-Pannier, L. (2014). The early development of brain white matter: a review of imaging studies in fetuses, newborns and infants. neuroscience, 276, 48-71.

Dubois, J., Adibpour, P., Poupon, C., Hertz-Pannier, L., & Dehaene-Lambertz, G. (2016). MRI and M/EEG studies of the white matter development in human fetuses and infants: review and opinion. Brain Plasticity, 2(*1*), 49-69.

Dubois, J., Lefèvre, J., Angleys, H., Leroy, F., Fischer, C., Lebenberg, J., … & Germanaud, D. (2019). The dynamics of cortical folding waves and prematurity-related deviations revealed by spatial and spectral analysis of gyrification. Neuroimage, 185, 934-946.

Dupont, E., Hanganu, I. L., Kilb, W., Hirsch, S., & Luhmann, H. J. (2006). Rapid developmental switch in the mechanisms driving early cortical columnar networks. Nature, 439(*7072*), 79-83.

Engelhardt, E., Inder, T. E., Alexopoulos, D., Dierker, D. L., Hill, J., Van Essen, D., & Neil, J. J. (2015). Regional impairments of cortical folding in premature infants. Annals of neurology, 77(*1*), 154-162.

Fenn-Moltu, S., Fitzgibbon, S. P., Ciarrusta, J., Eyre, M., Cordero-Grande, L., Chew, A., … & Batalle, D. (2023). Development of neonatal brain functional centrality and alterations associated with preterm birth. Cerebral Cortex, 33(*9*), 5585-5596.

França, L. G., Ciarrusta, J., Gale-Grant, O., Fenn-Moltu, S., Fitzgibbon, S., Chew, A., … & Batalle, D. (2024). Neonatal brain dynamic functional connectivity in term and preterm infants and its association with early childhood neurodevelopment. Nature Communications, 15(1), 16.

Gao, W., Alcauter, S., Elton, A., Hernandez-Castillo, C. R., Smith, J. K., Ramirez, J., & Lin, W. (2015). Functional network development during the first year: relative sequence and socioeconomic correlations. Cerebral cortex, 25(*9*), 2919-2928.

Gondová, A., Neumane, S., Leprince, Y., Mangin, J. F., Arichi, T., & Dubois, J. (2023). Predicting neurodevelopmental outcomes from neonatal cortical microstructure: A conceptual replication study. Neuroimage: Reports, 3(*2*), 100170.

Gozdas, E., Parikh, N. A., Merhar, S. L., Tkach, J. A., He, L., & Holland, S. K. (2018). Altered functional network connectivity in preterm infants: antecedents of cognitive and motor impairments? Brain Structure and Function, 223, 3665-3680.

Grigg-Damberger, M. M. (2016). The visual scoring of sleep in infants 0 to 2 months of age. Journal of clinical sleep medicine, 12(3), 429–445.

Gui, A., Bussu, G., Tye, C., Elsabbagh, M., Pasco, G., Charman, T., … & Jones, E. J. (2021). Attentive brain states in infants with and without later autism. Translational psychiatry, 11(*1*), 196.

Guyer, C., Werner, H., Wehrle, F., Bölsterli, B. K., Hagmann, C., Jenni, O. G., & Huber, R. (2019). Brain maturation in the first 3 months of life, measured by electroencephalogram: A comparison between preterm and term-born infants. Clinical Neurophysiology, 130(*10*), 1859-1868.

Hermans, T., Khazaei, M., Raeisi, K., Croce, P., Tamburro, G., Dereymaeker, A., … & Comani, S. (2023). Microstate Analysis reflects maturation of the Preterm Brain. Brain Topography, 1-14.

Hintz, S. R., Kendrick, D. E., Vohr, B. R., Poole, W. K., Higgins, R. D., & Nichd Neonatal Research Network. (2006). Gender differences in neurodevelopmental outcomes among extremely preterm, extremely-low-birthweight infants. Acta paediatrica, 95(*10*), 1239-1248.

Kelly, C. E., Thompson, D. K., Adamson, C. L., Ball, G., Dhollander, T., Beare, R., … & Inder, T. E. (2023). Cortical growth from infancy to adolescence in preterm and term-born children. *Brain*, awad348.

Kelly, L. A., Branagan, A., Semova, G., & Molloy, E. J. (2023). Sex differences in neonatal brain injury and inflammation. Frontiers in Immunology, 14, 1243364.

Kelly, C. E., Thompson, D. K., Adamson, C. L., Ball, G., Dhollander, T., Beare, R., … & Inder, T. E. (2023). Cortical growth from infancy to adolescence in preterm and term-born children. Brain, awad 348.

Keunen, K., Counsell, S. J., & Benders, M. J. (2017). The emergence of functional architecture during early brain development. Neuroimage, 160, 2–14.

Khazaei, M., Raeisi, K., Croce, P., Tamburro, G., Tokariev, A., Vanhatalo, S., … & Comani, S. (2021). Characterization of the Functional Dynamics in the Neonatal Brain during REM and NREM Sleep States by means of Microstate Analysis. Brain topography, 34, 555-567.

Kidokoro, H., Neil, J. J., & Inder, T. E. (2013). New MR imaging assessment tool to define brain abnormalities in very preterm infants at term. American Journal of Neuroradiology, 34(11), 2208–2214.

Koenig, T., Prichep, L., Lehmann, D., Sosa, P. V., Braeker, E., Kleinlogel, H., … & John, E. R. (2002). Millisecond by millisecond, year by year: normative EEG microstates and developmental stages. Neuroimage, 16(1), 41–48.

Kostović, & Judaš. (2015). Embryonic and fetal development of the human cerebral cortex. Brain Mapp, 2, 167–175.

Kostović, I., Sedmak, G., Vukšić, M., & Judaš, M. (2015). The relevance of human fetal subplate zone for developmental neuropathology of neuronal migration disorders and cortical dysplasia. CNS neuroscience & therapeutics, 21(*2*), 74-82.

Lehmann, D., Ozaki, H., & Pál, I. (1987). EEG alpha map series: brain micro-states by space-oriented adaptive segmentation. Electroencephalography and clinical neurophysiology, 67(3), 271–288

Li W, et al. (2017). REM sleep selectively prunes and maintains new synapses in development and learning. Nature Neuroscience

Liu, J., Xu, J., Zou, G., He, Y., Zou, Q., & Gao, J. H. (2020). Reliability and individual specificity of EEG microstate characteristics. Brain Topography, 33, 438–449.

Livingston, L. A., & Happé, F. (2017). Conceptualising compensation in neurodevelopmental disorders: Reflections from autism spectrum disorder. Neuroscience & Biobehavioral Reviews, 80, 729–742.

Maitre, N. L., Key, A. P., Chorna, O. D., Slaughter, J. C., Matusz, P. J., Wallace, M. T., & Murray, M. M. (2017). The dual nature of early-life experience on somatosensory processing in the human infant brain. Current Biology, 27(7), 1048–1054.

Marlow, N., Wolke, D., Bracewell, M. A., & Samara, M. (2005). Neurologic and developmental disability at six years of age after extremely preterm birth. New England journal of medicine, 352(1), 9–19.

Michel, C. M., & Koenig, T. (2018). EEG microstates as a tool for studying the temporal dynamics of whole-brain neuronal networks: a review. Neuroimage, 180, 577–593.

Murray, M. M., Brunet, D., and Michel, C. M. (2008). Topographic ERP analyses: A step-by-step tutorial review. Brain Topography, 20(4):249–264.

Neumane, S., Gondova, A., Leprince, Y., Hertz-Pannier, L., Arichi, T., & Dubois, J. (2022). Early structural connectivity within the sensorimotor network: Deviations related to prematurity and association to neurodevelopmental outcome. Frontiers in Neuroscience, 16, 932386’

O’Driscoll, D. N., McGovern, M., Greene, C. M., & Molloy, E. J. (2018). Gender disparities in preterm neonatal outcomes. Acta Paediatrica, 107(*9*), 1494-1499.

Omidvarnia, A., Fransson, P., Metsäranta, M., & Vanhatalo, S. (2014). Functional bimodality in the brain networks of preterm and term human newborns. Cerebral cortex, 24(*10*), 2657-2668.

Périvier, M., Rozé, J. C., Gascoin, G., Hanf, M., Branger, B., Rouger, V., … & Flamant, C. (2016). Neonatal EEG and neurodevelopmental outcome in preterm infants born before 32 weeks. Archives of Disease in Childhood-Fetal and Neonatal Edition, 101(3), F253-F259.

Pierrat, V., Marchand-Martin, L., Arnaud, C., Kaminski, M., Resche-Rigon, M., Lebeaux, C., … & EPIPAGE-2 Writing Group. (2017). Neurodevelopmental outcome at 2 years for preterm children born at 22 to 34 weeks’ gestation in France in 2011: EPIPAGE-2 cohort study. bmj, 358.

Pierrat, V., Marchand-Martin, L., Marret, S., Arnaud, C., Benhammou, V., Cambonie, G., … & Ancel, P. Y. (2021). Neurodevelopmental outcomes at age 5 among children born preterm: EPIPAGE-2 cohort study. bmj, 373.

Poulsen, A. T., Pedroni, A., Langer, N., & Hansen, L. K. (2018). Microstate EEGlab toolbox: An introductory guide. bioRxiv, 289850.

Rupawala, M., Bucsea, O., Laudiano-Dray, M. P., Whitehead, K., Meek, J., Fitzgerald, M., … & Fabrizi, L. (2023). A developmental shift in habituation to pain in human neonates. Current Biology, 33(*8*), 1397-1406.

Takarae, Y., Zanesco, A., Keehn, B., Chukoskie, L., Müller, R. A., & Townsend, J. (2022). EEG microstates suggest atypical resting-state network activity in high-functioning children and adolescents with autism spectrum development. Developmental science, 25(*4*), e13231.

Thompson, D. K., Kelly, C. E., Chen, J., Beare, R., Alexander, B., Seal, M. L., … & Spittle, A. J. (2019). Characterisation of brain volume and microstructure at term-equivalent age in infants born across the gestational age spectrum. NeuroImage: Clinical, 21, 101630.

Tokariev, A., Roberts, J. A., Zalesky, A., Zhao, X., Vanhatalo, S., Breakspear, M., & Cocchi, L. (2019). Large-scale brain modes reorganize between infant sleep states and carry prognostic information for preterms. Nature communications, 10(*1*), 2619.

Wehrle, F. M., Michels, L., Guggenberger, R., Huber, R., Latal, B.’ O’Gorman, R. L., & Hagmann, C. F. (2018). Altered resting-state functional connectivity in children and adolescents born very preterm short title. NeuroImage: Clinical, 20, 1148-1156.

Whitehead, K., Pressler, R., & Fabrizi, L. (2017). Characteristics and clinical significance of delta brushes in the EEG of premature infants. Clinical neurophysiology practice, 2, 12-18.

World Health Organisation (WHO) 2024 preterm birth fact sheet. Available at: https://www.who.int/news-room/fact-sheets/detail/preterm-birth

Wu, D., Chang, L., Akazawa, K., Oishi, K., Skranes, J., Ernst, T., & Oishi, K. (2017). Mapping the critical gestational age at birth that alters brain development in preterm-born infants using multi-modal MRI. Neuroimage, 149, 33-43.

Wynn, J. L., Wong, H. R., Shanley, T. P., Bizzarro, M. J., Saiman, L., & Polin, R. A. (2014). Time for a neonatal-specific consensus definition for sepsis. Pediatric Critical Care Medicine, 15(*6*), 523-528.

Yrjölä, P., Stjerna, S., Palva, J. M., Vanhatalo, S., & Tokariev, A. (2022). Phase-based cortical synchrony is affected by prematurity. Cerebral Cortex, 32(10), 2265–2276.

Yakovlev P. L. & Lecours, A. R. in Regional Development of the Brain in Early Life (ed. Minkowski, A.) 3–69 (Blackwell, Oxford, 1967).

Zhang, K., Shi, W., Wang, C., Li, Y., Liu, Z., Liu, T., … & Wang, G. (2021). Reliability of EEG microstate analysis at different electrode densities during propofol-induced transitions of brain states. NeuroImage, 231, *Article 117861*.

## References

Antonova, E., Holding, M., Suen, H. C., Sumich, A., Maex, R., & Nehaniv, C. (2022). EEG microstates: functional significance and short-term test-retest reliability. Neuroimage: Reports, 2(2), 100089

Kleinert, T., Koenig, T., Nash, K., & Wascher, E. (2024). On the reliability of the EEG microstate approach. Brain topography, 37(2), 271–286.

Jun, S., Alderson, T. H., Malone, S. M., Harper, J., Hunt, R. H., Thomas, K. M., … & Sadaghiani, S. (2024). Rapid dynamics of electrophysiological connectome states are heritable. Network Neuroscience, 1–50.

